# An orthogonal differentiation platform for genomically programming stem cells, organoids, and bioprinted tissues

**DOI:** 10.1101/2020.07.11.198671

**Authors:** Mark A. Skylar-Scott, Jeremy Y. Huang, Aric Lu, Alex H.M. Ng, Tomoya Duenki, Lucy L. Nam, Sarita Damaraju, George M. Church, Jennifer A. Lewis

## Abstract

Simultaneous differentiation of human induced pluripotent stem cells (hiPSCs) into divergent cell types offers a pathway to achieving tailorable cellular complexity, patterned architecture, and function in engineered human organoids and tissues. Recent transcription factor (TF) overexpression protocols typically produce only one cell type of interest rather than the multitude of cell types and structural organization found in native human tissues. Here, we report an orthogonal differentiation platform for genomically programming stem cells, organoids and bioprinted tissues with controlled composition and organization. To demonstrate this platform, we orthogonally differentiated endothelial cells and neurons from hiPSCs in a one-pot system containing neural stem cell-specifying media. By aggregating inducible-TF and wildtype hiPSCs into pooled and multicore-shell embryoid bodies, we produced vascularized and patterned cortical organoids within days. Using multimaterial 3D bioprinting, we patterned 3D neural tissues from densely cellular, matrix-free stem cell inks that were orthogonally differentiated on demand into distinct layered regions composed of neural stem cells, endothelium, and neurons, respectively. Given the high proliferative capacity and patient-specificity of hiPSCs, our platform provides a facile route for programming cells and multicellular tissues for drug screening and therapeutic applications.

## Main

Recent innovations in organoid development and 3D bioprinting offer emerging pathways to creating autologous human tissues for drug screening and therapeutic applications^1–4^. The ability to generate organoids via self-assembly and differentiation of embryoid bodies (EBs), aggregated from hiPSCs, offers a bottom-up approach to organ-like microarchitectures^5,6^. By contrast, multimaterial 3D bioprinting offers a top-down method for fabricating heterogeneous stem-cell derived tissues^7–11^. However, both techniques are limited by the speed, efficiency, and scalability of stem cell differentiation. While protocols for generating cerebral^12–14^, renal^15–17^, retinal^18^ and other organoids have recently been reported, they often rely on prolonged culture times ranging from weeks to several months to approach organ-level cellular diversity. Furthermore, organoid protocols that generate a wider range of cells and tissues generally result in less reproducible organoids, resulting in a trade-off between organoid reproducibility and cellular diversity^19,20^. A similarly large number of cell types, each differentiated and rendered into densely cellular bioinks, would be required for multimaterial 3D bioprinting of bulk organ-specific tissues. Hence, new approaches that enhance the efficiency, speed, scalability, and tailorability of stem cell differentiation are needed to generate programmable multicellular organoids and tissues from pluripotent stem cells.

To guide stem cell differentiation, one can introduce extracellular cues by controlling the media composition or modulate their intracellular state by overexpressing various transcription factors (TFs). Many TF-based protocols synergistically apply external and internal cues to promote rapid and efficient cell differentiation to a single lineage; examples include the (1) derivation of endothelial cells cultured in endothelial cell growth medium, while overexpressing ETS translocation variant 2 (ETV2)^21,22^, (2) derivation of neurons cultured in neurobasal-A medium with B27, while overexpressing neurogenin-1 and −2 (NGN1, NGN2)^23^, and (3) derivation of hepatocytes cultured in hepatocyte medium, while overexpressing combinations of Gata4, Hnf1α, Hnf4α, Foxa1, Foxa2, and Foxa3^24,25^. While these approaches aim to differentiate cells into a singular phenotype, human tissues are composed of multiple cell types organized into hierarchically patterned structures. While the overexpression of ‘less-specific’ TFs may create progenitors for multiple cell types^26^, this strategy offers limited control over the precise composition and distribution of cell types in the resulting tissue. As an alternate approach, one can overexpress TFs in a subset of cells within a developing organoid to guide their differentiation into selected and precise phenotypes. For example, Cakir *et al.* recently combined TF-driven differentiation with traditional cortical organoid culture by inducing overexpression of ETV2 in a subset of cells within cortical organoids, giving rise to a vascular endothelium akin to brain microvasculature^27^. However, the generation of more complex multicellular tissues requires not only a broader range of programmable cell types, but methods that simultaneously enable control over their spatial patterning.

We posit that an ideal method for generating programmable multicellular organoids and 3D organ-specific tissues would begin by overexpressing multiple TFs, each of which is orthogonally differentiated to a specific cell type of interest with high efficiency. Next, by pooling or printing populations of wild type (WT) and engineered cell lines, one can programmably generate multicellular organoids and 3D tissues, respectively. Successful implementation would require the ability to simultaneously differentiate multiple hiPSC lines in a one-pot protocol independent of external cues provided by the cell culture media. Here, we report an orthogonal differentiation (OD) platform for achieving genomically programming human stem cells, organoids, and 3D bioprinted organ-specific tissues. We show that forced overexpression of intracellular TFs can operate independently of media-driven differentiation to generate specific cell types, each with near unity efficiency (**Fig. 1a**). When applied to randomly pooled or multicore-shell EBs, OD can be used to construct multicellular and spatially patterned organoids (**Fig. 1b**). Moreover, when OD is coupled with multimaterial 3D bioprinting of densely cellular, matrix-free WT and inducible-TF iPSC inks, pluripotent tissues can be patterned and subsequently transformed *in situ* to multicellular tissue constructs that mimic native tissue architectures (**Fig. 1c**). Although our OD platform can be broadly applied, we demonstrate its utility by creating vascularized cortical organoids in pooled and multicore-shell motifs as well as 3D cortical tissues composed of multiple cell types patterned in spatially distinct regions.

**Fig. 1.**
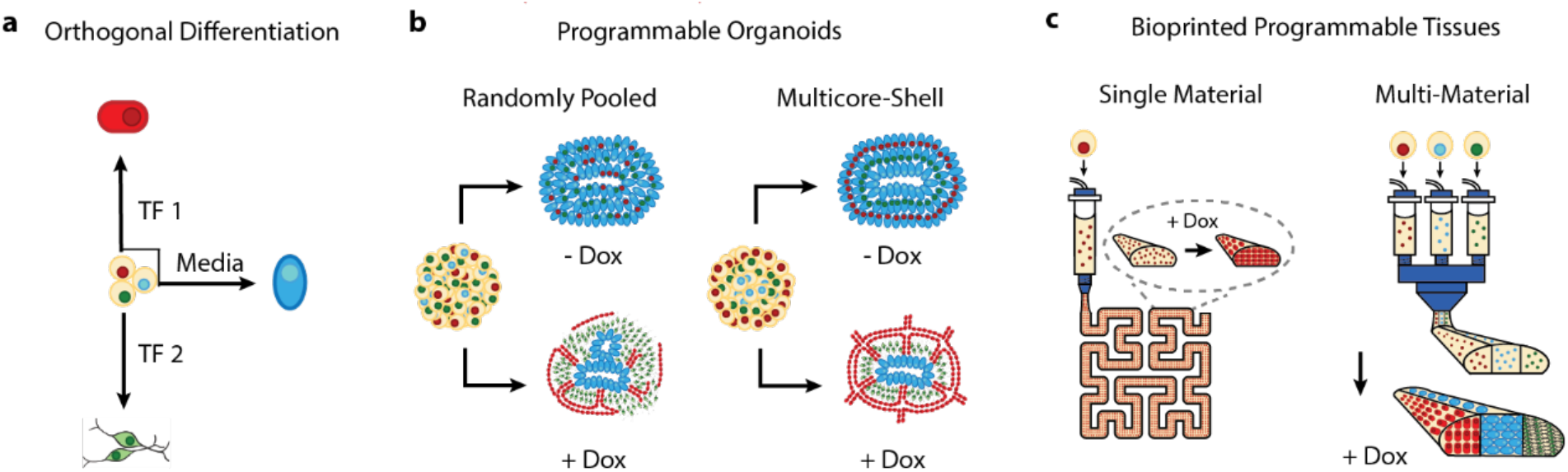
An orthogonal differentiation platform for genomically programming stem cells, organoids, and 3D bioprinted organ-specific tissues. **a**, Transcription factor overexpression generates distinct cell lineages with near-unity efficiency in identical cell media. **b**, Programmable and patterned organoids can be generated from randomly pooled and multicore-shell embryoid bodies via orthogonal differentiation. **c**, 3D organ-specific tissues composed of programmable pluripotent cell lines can be bioprinted and transformed *in situ* via orthogonal differentiation.

## Results

### Programming stem cell differentiation

The derivation of neural tissue from hiPSCs serves as an ideal example to assess the feasibility of orthogonal differentiation, given that pluripotent cells are efficiently differentiated into neuroectoderm in a dual-SMAD inhibiting medium^28^ (**Fig. 2**). We tested three different Personal Genome Project 1 (PGP1) hiPSC lines, one WT line and two inducible-TF lines, in identical medium conditions. WT PGP1 cells form neural stem cells when cultured in neural induction medium (NIM) containing TGF-ß- and BMP4-pathway small molecule inhibitors, as previously reported^29^ (**Fig. 2a**). For the inducible-TF lines, we utilized a PiggyBac transposon vector to incorporate an all-in-one Tet-On system that enables the rapid and highly efficient doxycycline-induced upregulation of transcription factors. To generate inducible endothelial (iEndo) cells, we overexpressed the transcription factor ETV2, which is known to drive rapid and efficient directed differentiation of pluripotent cells to vascular endothelial cells^22^ (**Fig. 2b**). To generate inducible neurons (iNeuron), we upregulated NGN1, as single or co-expression of neurogenins, which is known to rapidly generate neurons from stem cells^23,30^ (**Fig. 2c**).

**Fig. 2.**
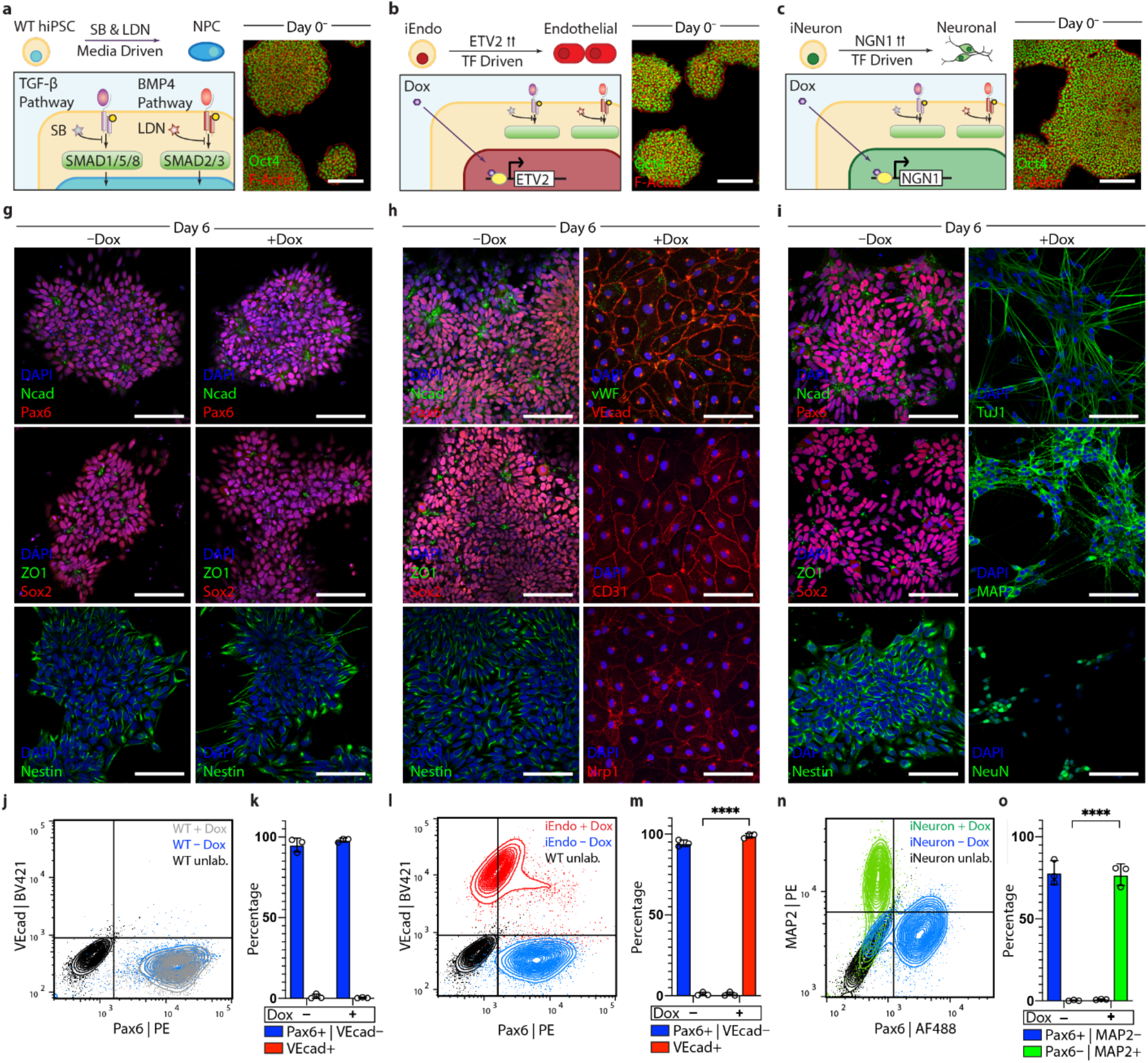
Programmable differentiation of pluripotent stem cells via orthogonal differentiation under identical media conditions. **a**, Left, schematic detailing WT PGP1 hiPSC differentiation into neural stem cells in NIM. Right, immunostaining of Oct4 and F-actin of WT PGP1 colonies on day 0. **b**, Left, schematic detailing iEndo differentiation through doxycycline-induced ETV2 isoform-2 overexpression. Right, immunostaining of Oct4 and F-actin of iEndo colonies on day 0. **c**, Left, schematic detailing iNeuron differentiation through doxycycline-induced NGN1 overexpression in culture. Right, immunostaining of Oct4 and F-actin of iNeuron colonies on day 0. **d**, WT PGP1s cultured in NIM for 6 days without (left column) or with (right column) doxycycline. Immunostaining of N-cadherin and Pax6, ZO1 and Sox2, and nestin. **e**, iEndo cells cultured in NIM for 6 days without (left column) or with (right column) doxycycline. Left, immunostaining of N-cadherin and Pax6, ZO1 and Sox2, and nestin. Right, immunostaining of vWF and VE-cadherin, CD31, and Nrp1. **f**, iNeuron cells cultured in NIM for 6 days without (left column) or with (right column) doxycycline. Left, immunostaining of N-cadherin and Pax6, ZO1 and Sox2, and nestin. Right, immunostaining of TuJ1, MAP2, and NeuN. **g**, Left, flow cytometry plots quantifying wild-type PGP1 differentiation to Pax6+ neural stem cells by 6 days with (grey) and without (blue) doxycycline. Right, quantification of populations. **h**, Left, flow cytometry plots indicating that in the presence of doxycycline iEndo PGP1s differentiate into VECad+ endothelium (red). In the absence of doxycycline, iEndo PGP1s differentiate into Pax6+ neural stem cells (blue). Right, quantification of populations, mean ± s.e.m., n = 3 biological replicates, **** P = 7.06×10^−7^, unpaired two-tailed t-test. **i**, Left, flow cytometry plots indicating that in the presence of doxycycline iNeuron PGP1s differentiate into MAP2+ neurons (green). In the absence of doxycycline, iNeurons differentiate into Pax6+ neural stem cells (blue). Right, quantification of populations, mean ± s.e.m., n = 3 biological replicates, **** P = 3.53×10^−5^, unpaired two-tailed t-test. Scale bars: 50 μm in **a-f**.

We characterized the differentiation of the WT and inducible-TF hiPSC lines in the absence (control) and presence of doxycycline. When cultured for six days in NIM, WT hiPSC cells experienced a loss of pluripotency and efficiently differentiated into a neural stem cell fate in both cases, as indicated by the formation of characteristic polarized neural rosettes and the expression of neural cadherin (Ncad), paired-box gene 6 (Pax6), SRY-box transcription factor 2 (Sox2) and Nestin (**Fig. 2d**). Similarly, iEndo cells undergo media-driven differentiation to neural stem cells in the absence of doxycycline (**Fig. 2e**). However, in the presence of doxycycline, the overexpression of ETV2 drives the rapid differentiation to a vascular endothelial cell phenotype, as evidenced by the formation of a confluent cobblestone morphology, the expression of vascular endothelial cadherin (VECad), von-Willebrand factor (vWF), CD31, and the vascular endothelial growth factor receptor Neuropilin 1 (NRP1). iNeuron cells also differentiated into neural stem cells in the absence of doxycycline. In the presence of doxycycline, iNeurons formed neurons with a bipolar morphology and expressed neural markers Tuj1, microtubule associated protein 2 (MAP2) and neuronal nuclei (NeuN) (**Fig. 2f**). When induced, both iEndo and iNeuron efficiently and rapidly differentiated, confirming the feasibility of OD (**Fig. 2g-i, Fig. S1**). Indeed, the stark contrast between the differentiation of iEndo in the presence (99% vascular endothelium) and absence (94% neural stem cells) of doxycycline serves as a striking illustration of the orthogonality of external (media-driven) versus internal (TF-directed) cell differentiation. The data clearly demonstrate that intracellular TF forced differentiation can fully override the otherwise strong and specific extracellular dual-SMAD inhibition cues (**Fig. 2h**).

Next, we investigated whether simultaneous one-pot orthogonal differentiation of a mixed initial population of iPSCs would give rise to a heterogeneous differentiated cell population with programmed composition. Specifically, we seeded different proportions of WT, iEndo, and iNeuron hiPSCs onto a Matrigel surface and cultured them in NIM with doxycycline. We observed that distinct multicellular populations form after six days in culture (**Fig. S2**). Furthermore, under these conditions, iEndo cells self-assembled into a network-like microvasculature that is consistent with endothelial tubulogenesis assays. By contrast, WT cells formed neurospheres that rose above the surface of the gel, while iNeuron cells formed a network of neurites. Notably, when iEndo and WT cells are cultured together, the differentiated cells form a distinct network pattern in which the endothelial cells comprise the edges and the neurospheres comprise the nodes. Using iEndo-mKate2 and WT-GFP labeled cells, we confirmed that the vascular and neurosphere components are indeed composed of iEndo and WT cells, respectively (**Movie 1**). When iEndo and iNeuron cells are cultured together, the resulting endothelial and neuronal cells formed overlapping networks in which neurites extend along the vascular network. To demonstrate simultaneous OD, we pooled all three cell lines together and cultured in NIM and in the absence (control) and presence of doxycycline. Absent doxycycline, all three cell lines differentiated into neurospheres, with no visible VE-Cad or MAP2 staining. However, in the presence of doxycycline, WT, iEndo, and iNeuron cells are differentiated into Sox2+ neurospheres, VE-Cad+ vascular endothelium, and MAP2+ neurons, respectively. Importantly, our results suggest that orthogonal differentiation could be used to greatly enhance organoid differentiation protocols, which typically aim to derive cells from a single germ layer^12–14,19,20^.

### Programmable cortical organoids

Having demonstrated an OD protocol that is capable of near-unity efficiency of ETV2-driven differentiation of endothelium in NIM, we next sought to apply OD to address the challenge of organoid vascularization, since traditional cerebral (and cortical) organoid protocols fail to generate an embedded vascular network^12–14^. One naïve strategy to achieving this goal is to simply mix endothelial cells with hiPSCs to form a multicellular embryoid body that subsequently differentiates into the desired vascularized organoid. However, we find that when human umbilical vein endothelial cells (HUVECs) are combined with WT PGP1 hiPSCs in microwells they phase separate within 24 h (**Fig. S3a**). This observation arises due to differences in cell adhesion^31^, as endothelial cells and epithelial hiPSCs express different cell adhesion molecules. Importantly, when iEndo-mKate2 and WT-GFP hiPSCs are mixed and cultured in microwells, single embryoid bodies are formed with interspersed cells (**Fig S3b, Fig 3b-c**). By controlling the ratio of WT-to-iEndo cells seeded in each microwell, we can precisely tailor the resulting endothelium in the organoids after orthogonal differentiation (**Fig S3c-d**). To illustrate this, we produced vascularized cortical organoids by pooling 67% WT hiPSCs and 33% iEndo cells into embryoid bodies that are cultured in dual-SMAD inhibiting media to direct dorsal forebrain formation. After 3 days in suspension culture, the embryoid bodies are embedded into a Matrigel-collagen extracellular gel droplet. At 10 days, the avascular WT-only organoids (control) retained a smooth border, while the organoids comprised of 2:1 ratio of WT-to-iEndo cells formed extensive sprouted vascular networks that penetrate into the surrounding gel, as marked by lectin and visualized using bright-field (**Fig. 3d**) and fluorescence microscopy (**Fig. 3e**). The complete absence of microvascular markers in the WT organoids is consistent with previous reports that cortical organoids, in the absence of iEndo cells, lack a vascular network.

**Fig. 3.**
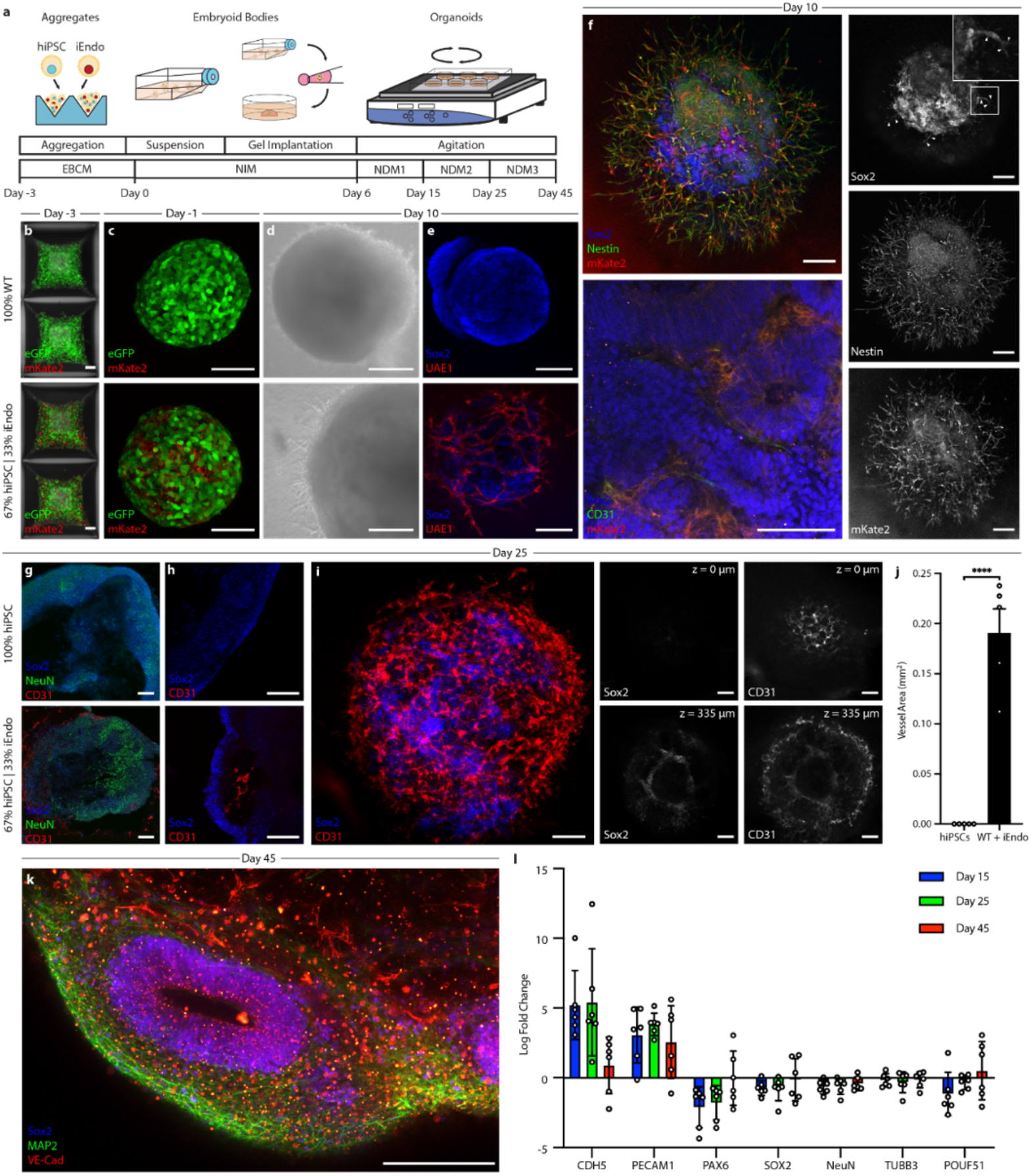
Programmable vascularization of cortical organoids via orthogonal differentiation. **a**, Schematic of the vascularized cortical organoid protocol. **b**, Fluorescent images of hiPSC aggregates in microwells three days before suspension culture. Top, 100% WT-eGFP PGP1 aggregates. Bottom, 67% WT-eGFP PGP1 and 33% iEndo-mKate2 randomly pooled cells in microwells. **c**, Fluorescent images of resulting EBs. Top, 100% WT-eGFP PGP1 EBs. Bottom, 67% WT-eGFP PGP1 and 33% iEndo-mKate2 randomly pooled EBs. **d**, Brightfield images of organoids derived from EBs cultured for 10 days. Top, 100% WT-eGFP PGP1 organoids. Bottom, 67% WT-eGFP PGP1 and 33% iEndo-mKate2 pooled organoids. **e**, Immunostaining of Sox2 and UAE1 of organoids cultured for 10 days. Top, 100% WT organoids. Bottom, 67% WT PGP1s and 33% iEndo pooled organoids. **f**, Left, top, confocal maximum intensity z-projections obtained from immunostaining of Sox2 and nestin of 67% WT PGP1s and 33% iEndo-mKate2 pooled organoids cultured for 10 days; right column, individual channels for Sox2, nestin, and mKate2. Arrowheads indicate Sox2+ positive cells that co-localize with the vasculature. Left, bottom, immunostaining of Sox2 and CD31 with iEndo-mKate2 of 67% WT PGP1s and 33% iEndo-mKate2 pooled organoids cultured for 10 days. **g**, Immunostaining of Sox2, NeuN, and CD31 of organoids cultured for 25 days. Top, 100% WT PGP1 organoids. Bottom, 67% WT PGP1 and 33% iEndo pooled organoids. **h**, Immunostaining of Sox2 and CD31 of organoids cultured for 25 days. Top, 100% WT PGP1 organoids. Bottom, 67% WT PGP1 and 33% iEndo pooled organoids. **i**, Left, maximum intensity z-projection with immunostaining of Sox2 and CD31 of 67% WT PGP1 and 33% iEndo pooled organoids cultured for 25 days. Right, optical slices of Sox2 and CD31 channels at depths of z = 0 μm and z = 335 μm. **j**, Quantification of vascular area in slices of a single 100% WT PGP1 organoid and a single 67% WT PGP1 and 33% iEndo pooled organoids. Data represents mean ± s.e.m. (n = 5, unpaired two-tailed t-test) **** P = 4.23×10^−5^. **k**, Maximum intensity z-projection with immunostaining of Sox2, MAP2, and VE-Cadherin of 67% WT PGP1 and 33% iEndo pooled organoids cultured for 45 days. **l**, RT-qPCR log fold-change expression bar-plot of 67% WT PGP1 and 33% iEndo pooled organoid versus 100% WT organoid. Data represents mean ± s.e.m. (n = 6, from three independent batches). Scale bars: 200 μm in **b**, **d**, **f**, **h**, **i**, and **k**; 100 μm in **c**, **e**, and **g**.

By pooling fluorescent mKate2-expressing iEndo cells, we confirmed that the embedded and sprouting vascular network arises solely from those cells, as evidenced by the co-localization of CD31 and mKate2 (**Fig. 3f**). Intriguingly, we see evidence of Sox2+ neural stem cells colocalizing along the sprouting vasculature (arrowheads, **Fig. 3f**). This interaction is consistent with our observations in co-culture of neural stem cells and iEndo endothelium on Matrigel hydrogels (**Fig. S2**) as well as known *in vivo* interactions of neural stem cells and endothelium in the neural stem cell niche in the subventricular zone^32^. Notably, endothelial cells reside in between the rosette or ventricle-like structures, whereas the Sox2+ germinal zone-like structures are devoid of these cells, consistent with observed vascular patterns in neurodevelopment^33^. After 25 days in culture, the vascularized cortical organoids developed NeuN positive neural populations (**Fig. 3g, Fig. S4**) and only organoids containing 33% iEndo cells possessed an embedded vascular network (**Fig. 3h**). Using whole-mount immunofluorescence of optically cleared organoids, we confirmed that this vascular network exists both at their surface and within their core (**Fig. 3i, Movie 2**). Embedded vascular networks are imaged in multiple slices within the cleared organoids, revealing a sharp difference between the WT-only and the WT+ iEndo cortical organoids (**Fig. 3j**). As expected, there is no detectable vasculature within the WT-only organoids. However, by day 45 of culture, WT+ iEndo cortical organoids developed ventricle-like architectures that are surrounded by neural cell bodies, neurites, and pervasive vascular network (**Fig. 3k, Fig. S5**). Using quantitative reverse transcription polymerase chain reaction (RT-qPCR), we find that these vascularized cortical organoids have significantly enhanced expression of vasculature-related genes (CDH5, PECAM1), while neural stem cell, neural, and pluripotent gene expression remain similar to the WT-only (control) organoids (**Fig. 3l**). This result demonstrates that orthogonal differentiation promotes vascular network formation without adversely affecting the neural phenotype within these organoids.

Moving beyond the random incorporation of iEndo cells into pooled embryoid bodies to form vascularized organoids, we then sought to create spatially patterned cortical organoids. We combined OD of WT, iEndo, and iNeuron cells in a multicore-shell organoid architecture that is reminiscent of a developing brain, i.e., a germinal zone (GZ) (inner core), surrounding neurons (middle layer), and an encapsulating perineural vascular plexus (outer shell) (**Fig. 4a**).

**Fig. 4.**
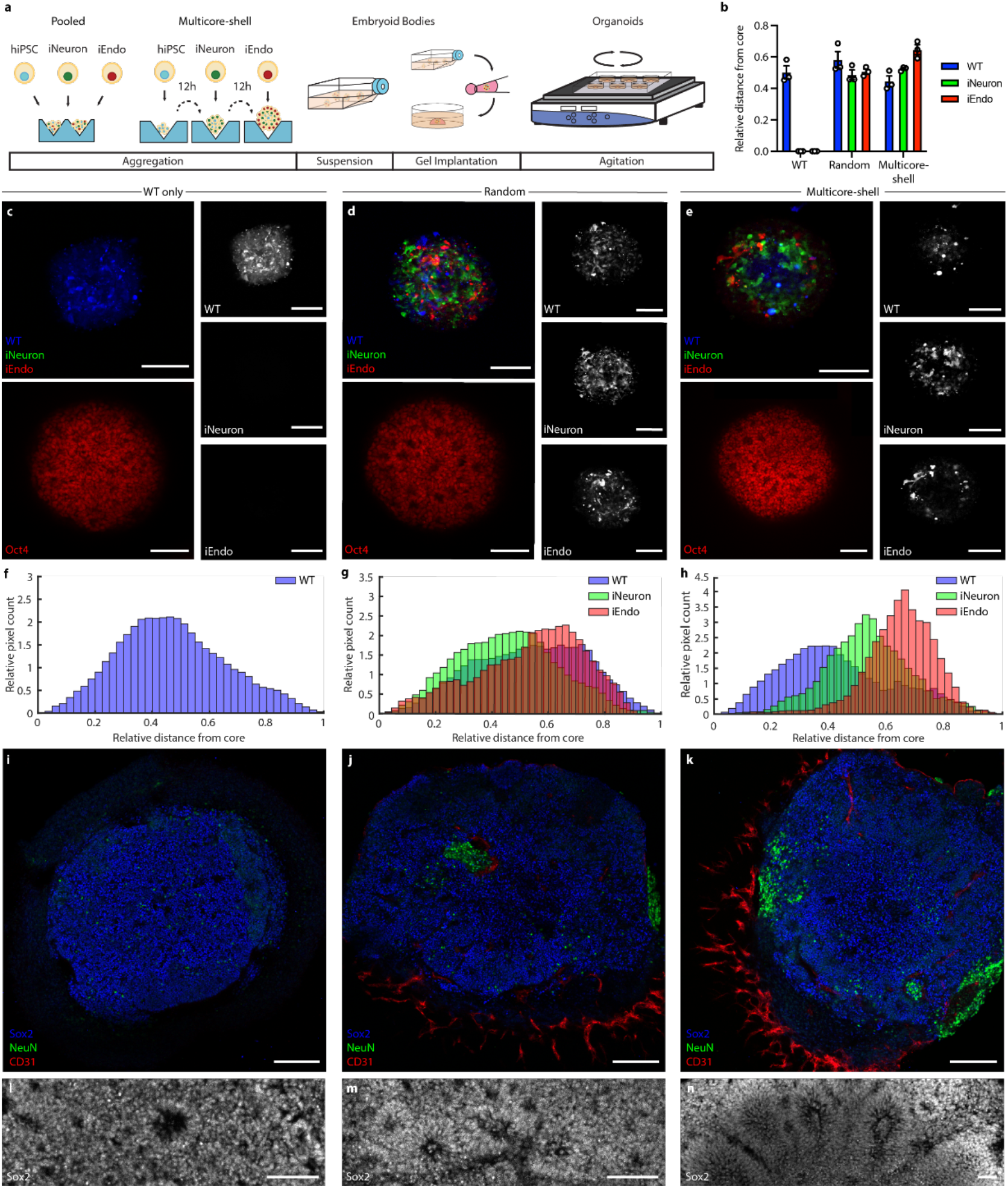
Multicore-shell cortical organoids. **a**, Schematic of the vascularized, multicore-shell organoid protocol. **b**, Quantification of hiPSCs seeding distribution of WT, iEndo, and iNeurons in WT-only, pooled, and multicore-shell approaches. Data represents the mean ± s.e.m. (n = 3, from three independent batches). **c**, WT-only organoids cultured for 1 day. Left, top, fluorescence image of CellTracker labeled WT-only organoids; right column, individual channels of CellTracker labeled WT, iNeuron, and iEndo cells. Left, bottom, immunostaining for Oct4. **d**, Randomly pooled organoids cultured for 1 day. Left, top, fluorescence image of CellTracker labeled randomly pooled organoids; right column, individual channels of CellTracker labeled WT, iNeuron, and iEndo cells. Left, bottom, immunostaining for Oct4. **e**, Multicore-shell organoids cultured for 1 day after final shell seeding. Left, top, fluorescence image of CellTracker labeled multicore-shell organoid; right column, individual channels of CellTracker labeled WT, iNeuron, and iEndo cells. Left, bottom, immunostaining for Oct4. Left-bottom, immunostaining for Oct4. **f**, Quantification of spatial distribution of cells in WT-only cortical organoids. **g**, Quantification of spatial distribution of cells in pooled cortical organoids. **h**, Quantification of spatial distribution of cells in multicore-shell cortical organoids. **i**, Immunostaining of Sox2, NeuN, and CD31 of WT-only organoids cultured for 10 days. **j**, Immunostaining of Sox2, NeuN, and CD31 of pooled organoids cultured for 10 days. **k**, Immunostaining of Sox2, NeuN, and CD31 of multicore-shell organoids cultured for 10 days. **l**, Immunostaining of Sox2 of WT organoids cultured for 10 days. **m**, Immunostaining of Sox2 of randomly pooled organoids cultured for 10 days. **n**, Immunostaining of Sox2 of multicore-shell organoids cultured for 10 days. Scale bars: 100 μm in **c-e**, **i-k**; 50 μm in **l-n**.

Additionally, the incorporation of iNeuron cells enables faster neurogenesis when compared to cortical organoids that typically require ~30 days of culture to form a NeuN+ neuronal population^34^. Stepwise addition of iPSCs in a U-well plate has previously been shown to generate a Janus embryoid body that can be used in conjunction with an inducible Sonic Hedgehog-expressing hiPSC line to encourage dorsoventral axis formation^35^. By using V-shaped wells, we created organoids with a radial-symmetric multicore-shell architectures, rather than two hemispherical compartments seen for Janus organoids assembled in U-wells (**Fig. 4b**). Embryoid bodies formed from WT-only, randomly pooled {WT, iNeuron, iEndo} triple populations, and step-wise multicore-shell organoids all formed cohesive Oct4+ embryoid bodies after one day in culture (**Fig. 4c-e**) with varying distributions of cell populations (**Fig. 4f-h**). Consistent with other cortical organoid protocols^34,36^, those containing WT-only cells contained few neurons at day 10, as evidenced by a minimal amount of NeuN expressing cells (**Fig. 4i**). By contrast, OD of iNeuron cells incorporated in random and multicore-shell organoids resulted in a large NeuN+ neuronal population by day 10. These neuronal populations surround the ventricle-like structures within the organoids, forming a neuron-rich layer that resides more superficially than the deeper GZs. We also find clusters of NeuN+ neurons deep in the center of randomly incorporated organoids that are absent in multicore-shell patterned organoids (**Fig. 4j-k**), suggesting that the initial spatial patterning imposed by the stepwise aggregation informs later layered organization within cortical organoids. Furthermore, a pervasive CD31+ vascular network is present throughout both randomly patterned and multicore-shell patterned organoids. Surprisingly, when compared to both the WT and randomly pooled embryoid bodies, orthogonal differentiation of patterned multicore-shell embryoid bodies gives rise to cortical organoids with larger ventricle-like structures with a more radially polarized architecture (**Fig. 4l-n**, **Fig. S6**).

### Programmable 3D neural tissues via bioprinting

Most bioprinting methods produce multicellular tissues by patterning inks composed of primary human cells suspended in hydrogel matrices. Recently, human mesenchymal stem cells (hMSCs)^37^ and hiPSC^38^ inks have been printed and differentiated *in situ* to form 3D tissue constructs. However, those stem-cell derived tissues were composed of single cell types defined by the differentiation protocol used. By leveraging the proliferative capacity of hiPSCs along with the programmability afforded by OD, single- and multicellular 3D human tissues can be created with high cell densities. As a simple demonstration, we first created concentrated, matrix-free hiPSC bioinks by pelleting single-cell suspensions of hiPSCs and then directly printing these pluripotent inks on transwell membranes (**Fig. 5a-c**, **Movie 3**). The printed filamentary features, which contain a remarkably high cellular density exceeding 5×10^8^ Oct4+ cells per ml (**Fig. 5d**), are subsequently encased in a gelatin-fibrinogen ECM to support the tissue and enable 3D cell migration. Importantly, calcein-AM/ethidium homodimer-2 Live/Dead assays of matrix-free hiPSC inks, printed through 50 μm and 100 μm tapered nozzles at speeds of up to 20 mm/s, confirm high cell viability akin to that observed for casted (control) tissues (**Fig. 5e**). The pluripotent filaments were printed with a high resolution, with widths measuring 132 μm and 182 μm (FWHM) for bioinks printed using 50 μm and 100 μm nozzles, respectively (**Fig. 5f**). Next, we printed planar patterns in the form of third-order pseudo-Hilbert curves composed of filamentary features produced from WT, iEndo, or iNeuron inks (**Fig. 5g-h**). Printed WT filaments exhibited a slight compaction when differentiated in NIM containing doxycycline, forming a neuroectoderm filament with small Ncad+ ventricle-like structures or rosettes punctuating its length. By contrast, printed and differentiated (iEndo) filaments exhibited vasculogenesis, resulting in the formation of a microvascular network over time. Finally, printed iNeuron filaments differentiated into densely packed neurons (NeuN+ cells) that formed a pervasive network of protruding Tuj1+ neurites.

**Fig. 5.**
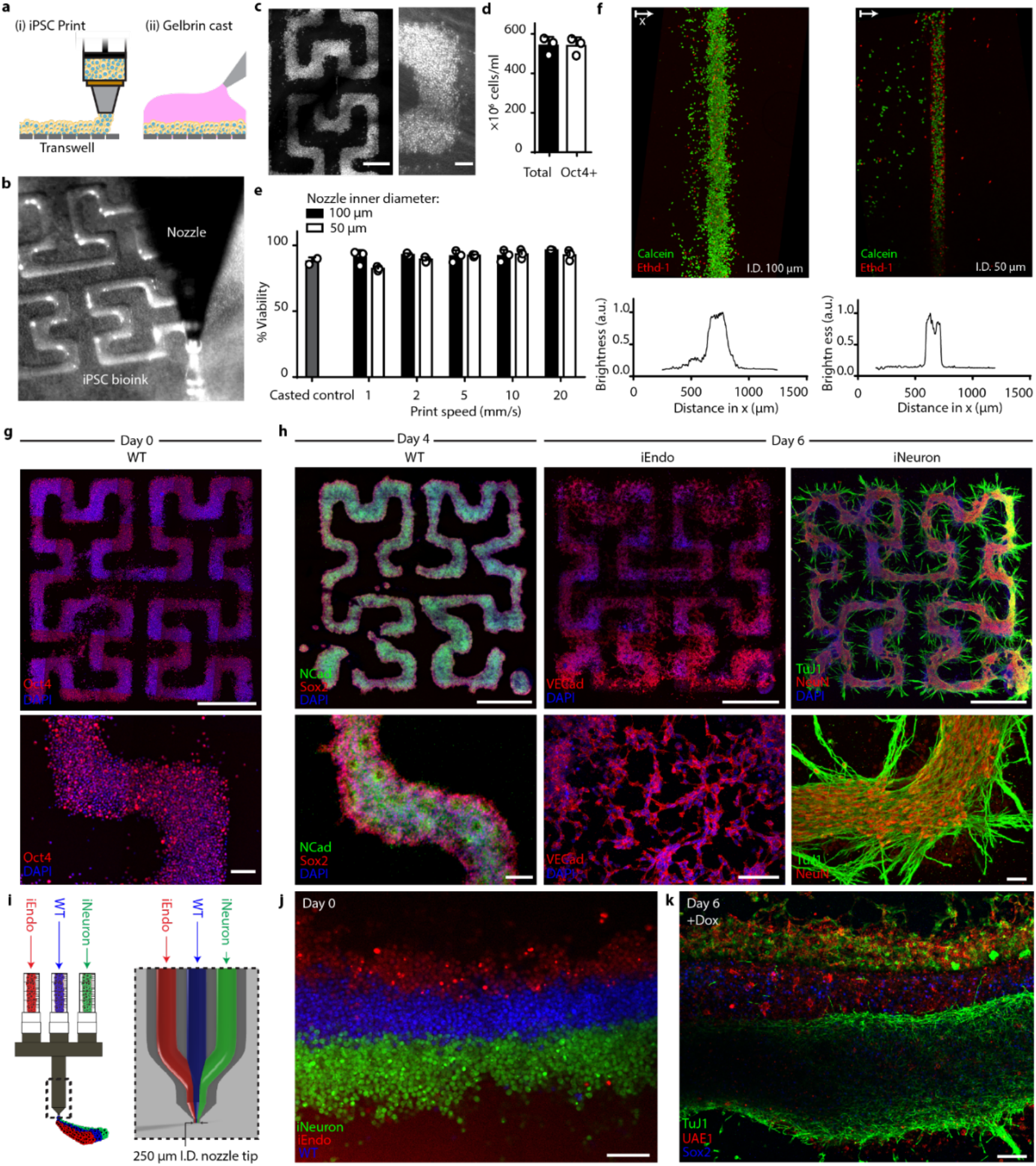
Multicellular neural tissues via 3D bioprinting coupled with orthogonal differentiation. **a**, Schematic of the bioprinting and gel casting process. **b**, Image of bioink extrusion through 50 μm nozzle during bioprinting process. **c**, Brightfield images of bioprinted filamentary features. **d**, Total cell number and flow-cytometry quantification of Oct4+ cells within bioprinted filaments immediately after printing. Data represents mean ± s.d. (n = 3 from three independent cell ink batches). **e**, hiPSC viability within bioprinted filaments at different speeds and casted (control) samples, as measured using calcein-AM/ethidium homodimer live/dead assays. Data represents mean ± s.e.m. (n = 3, from three independent batches). **f**, Live/dead staining of bioprinted filaments produced using nozzle diameters of 100 μm (top, left) and 50 μm (top, right) and the corresponding distributions (bottom) of their fluorescence intensity. **g**, Immunostaining of Oct4 in patterns fixed on day 0, immediately after printing. **h**, Bioprinted tissue architectures cultured in NIM. Left, immunostaining of N-cadherin and Sox2 for WT printed patterns at day 4. Middle, immunostaining of VE-cadherin for iEndo printed patterns at day 6. Right, immunostaining of TuJ1 and NeuN of iNeuron printed patterns at day 6. **i**, Design of triple-nozzle for multimaterial bioprinting. **j**, Fluorescence image of bioprinted CellTracker labeled WT, iEndo, and iNeuron inks. **k**, Immunostaining of TuJ1, UAE1 and Sox2 of 3D printed multicellular tissues at day 6. Scale bars: 500 μm in **g** (top) and **h** (top); 200 μm in **c** (left) and **i-k**; 50 μm in **c** (right), **g** (bottom), and **h** (bottom).

As a final demonstration, we created multicellular tissue architectures by co-printing three bioinks composed of WT, iEndo, and iNeuron hiPSCs, respectively (**Fig. 5i**, **Fig. S7**). The printhead contains three independent ink delivery channels that merge shortly upstream of the nozzle tip. Under laminar flow, these three inks converge to form tri-layer filaments that exit the nozzle. Using fluorescent dye-labeled WT, iEndo and iNeuron hiPSCs, we showed that this multimaterial printhead can successfully generate multicellular pluripotent filaments in layered architectures with high fidelity (**Fig. 5j**). After printing, we added doxycycline to induce the simultaneous orthogonal differentiation of neural stem cells, vascular endothelium, and neurons and observed that the layered architecture is preserved over 6 days of culture. (**Fig. 5k**). Finally, using this integrated OD and bioprinting platform, we recapitulated the geometry of a developing human dorsal forebrain coronal section by co-printing densely cellular WT and iNeuron hiPSC inks (**Fig. S8a-c**). After printing, we orthogonally differentiated the printed inks to form a neuroectoderm and overlying neuron-dense layer (**Fig. S8d**). As new protocols for deriving specific neuronal cell types emerge, our platform will enable patterning multilayer cortical architectures from an even broader array of inducible-TF hiPSCs.

## Discussion

We have demonstrated an orthogonal differentiation platform for rapidly programming and patterning human stem cells, organoids, and bioprinted organ-specific tissues within days. We found that overexpression of ETV2 and NGN1 efficiently overrides the dual SMAD-inhibiting media cues, enabling the simultaneous generation of vascular endothelium and neurons in a one-pot system. Leveraging this capability, we created both vascularized and multicore-shell brain organoids from pooled and patterned embryoid bodies, respectively, which contained larger, more distinct ventricle-like structures. Through multimaterial 3D bioprinting of matrix-free wildtype and inducible-TF hiPSC inks, we generated programmable, multicellular cortical tissues in layered architectures that can be orthogonally differentiated on demand. With further advancement of stem cell differentiation protocols, it may soon be possible to efficiently and scalably produce the multitude of cells present in the human body. Looking ahead, our platform offers a facile route for creating many types of programmable organoids and organ-specific tissues of interest for drug screening, disease modeling, and therapeutic applications.

---

**Fig. S1.**
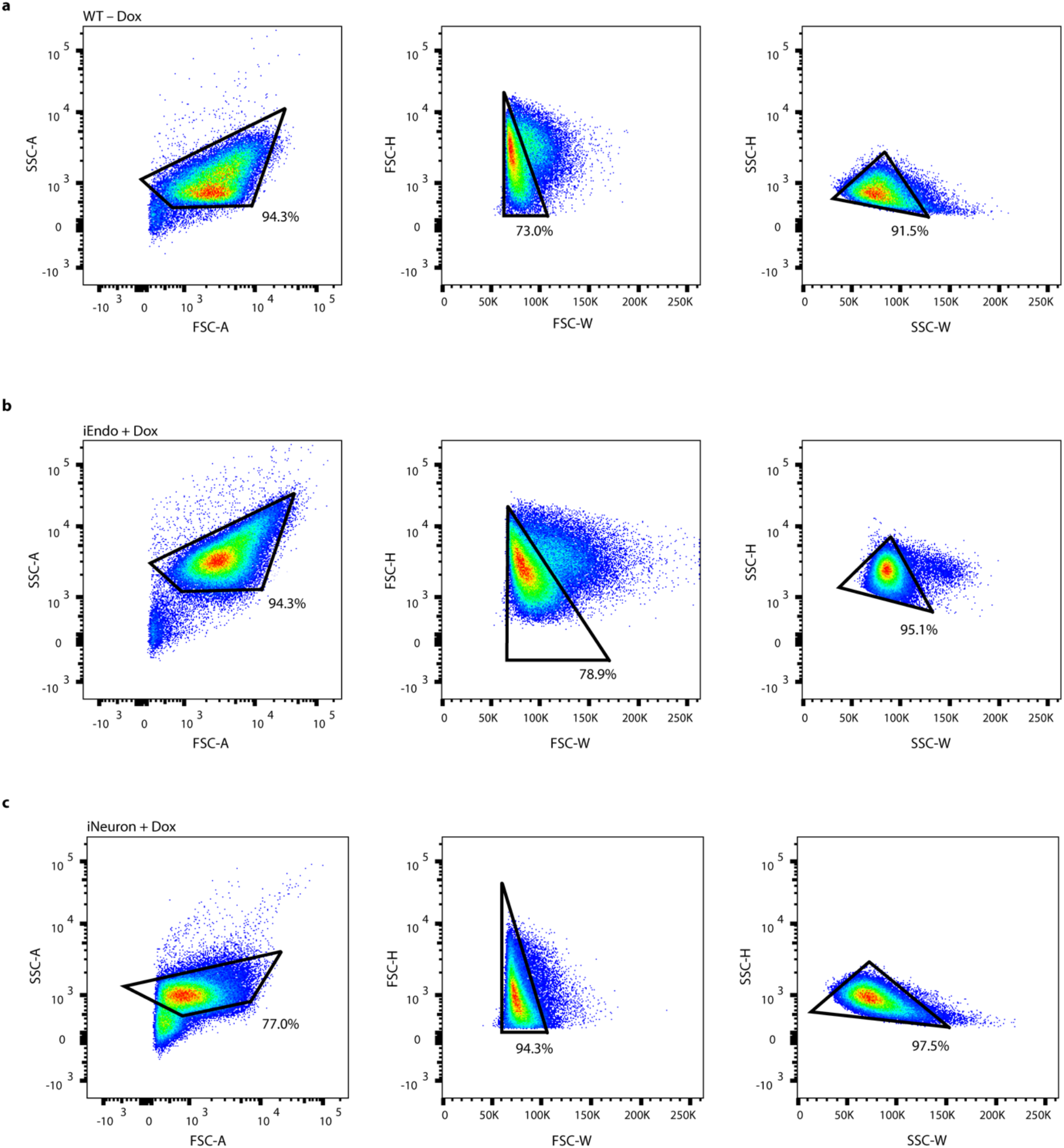
Flow cytometry gating examples. Gating strategies for **a** WT PGP1 cells, **b** iEndo cells, and **c** iNeuron cells. Left, forward-scatter (area) vs side-scatter (area). Middle, forward-scatter (width) vs forward-scatter (height). Right, side-scatter (width) vs side-scatter (height).

**Fig. S2.**
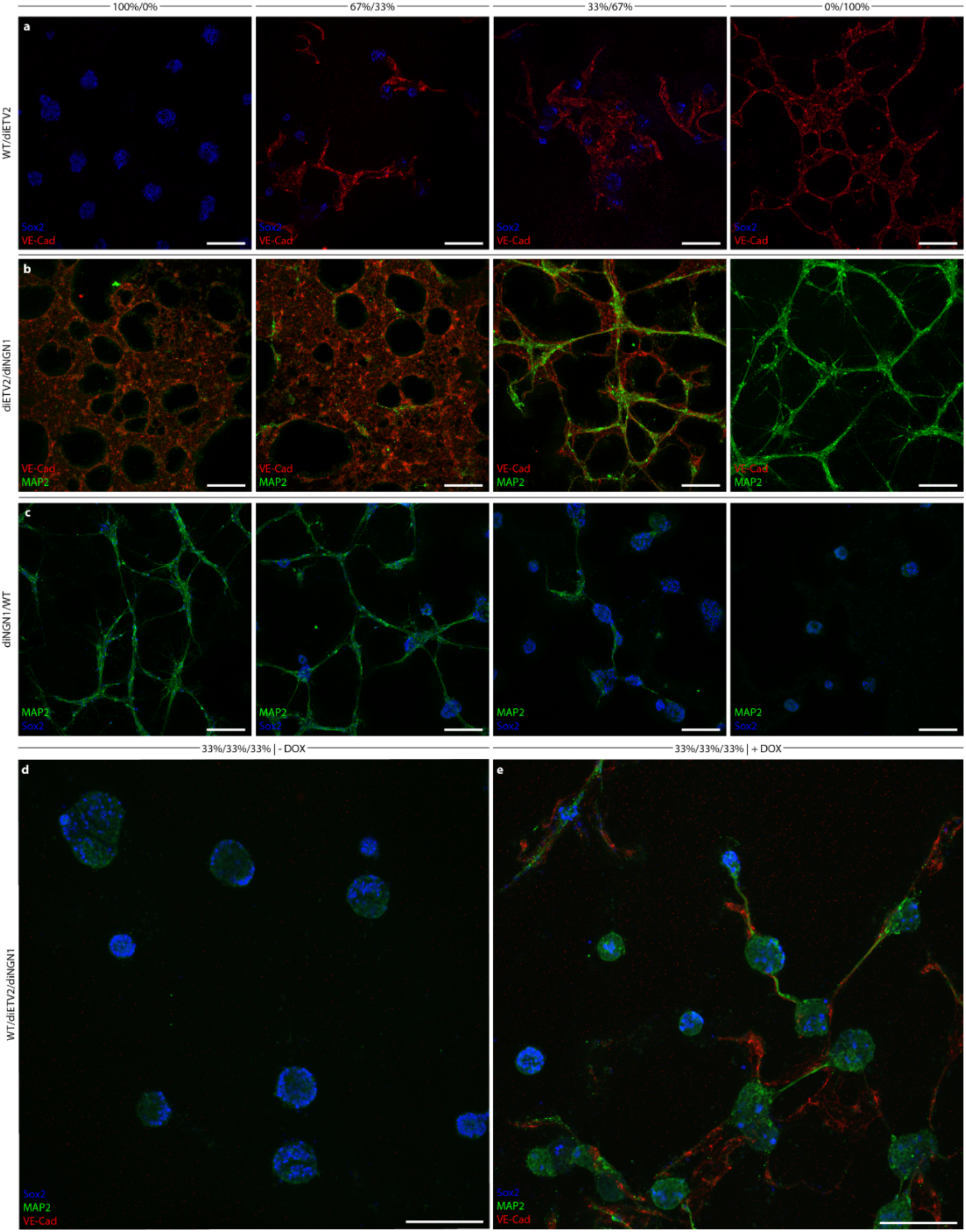
Orthogonal differentiation of pooled programmable iPSCs form tailorable multi-lineage cultures. **a**, Immunostaining of Sox2 and VE-Cadherin of different seeded proportions of WT PGP1 and iEndo cells on Matrigel hydrogels cultured in NIM for 6 days. Left, 100% WT culture. Middle-left, 67% WT and 33% iEndo. Middle-right, 33% WT and 67% iEndo. Right, 100% iEndo. **b**, Immunostaining of VE-Cadherin and MAP2 of different seeded proportions of iEndo and iNeuron on Matrigel hydrogels cultured in NIM for 6 days. Left, 100% iEndo. Middle-left, 67% iEndo and 33% iNeuron. Middle-right 33% iEndo and 67% iNeuron. Right, 100% iNeuron. **c**, Immunostaining of Map2 and Sox2 of different seeded proportions of WT PGP1s and iNeuron PGP1s on Matrigel hydrogels cultured in NIM for 6 days. Left, 100% iNeuron. Middle-left, 67% iNeuron and 33% WT. Middle-right 33% iNeuron and 67% WT. Right, 100% WT. **d**, Left, without doxycycline, immunostaining of Sox2, Map2, and VE-Cadherin of 33% WT, 33% iEndo, and 33% iNeuron on Matrigel hydrogel cultured in NIM for 6 days. Right, with doxycycline, immunostaining of Sox2, Map2, and VE-Cadherin of 33% WT, 33% iEndo, and 33% iNeuron on Matrigel hydrogels cultured in NIM for 6 days. Scale bars: 100 μm in **a-e**.

**Fig. S3.**
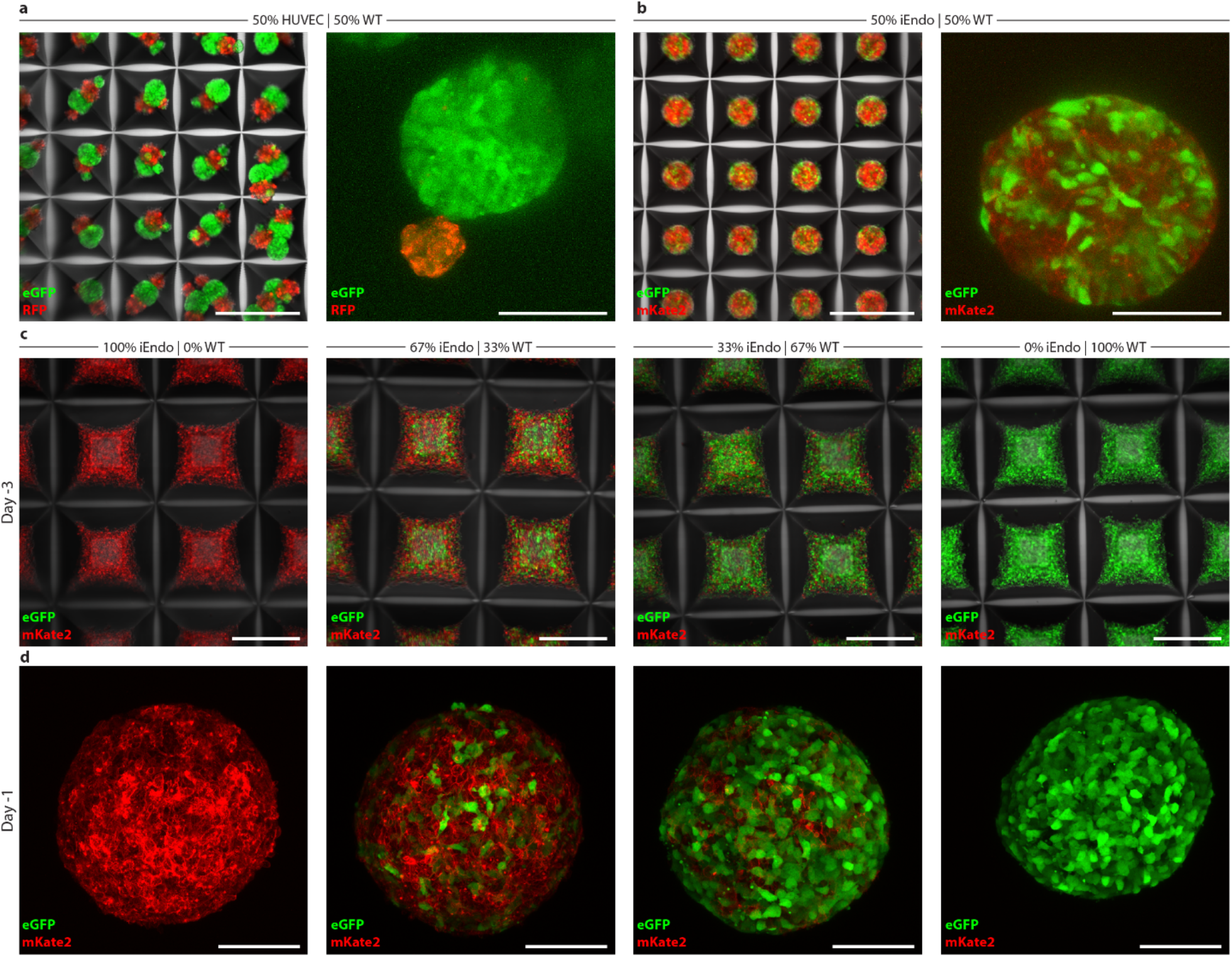
Pooled iPSCs enable the formation of cohesive embryoid bodies with tailorable cellular composition. **a**, Left, 50% WT-eGFP PGP1 and 50% RFP HUVEC EBs in microwell arrays cultured for 1 day. Right, 50% WT-eGFP PGP1 and 50% RFP HUVEC EBs cultured for 3 days. **b**, Left, 50% WT-eGFP PGP1 and 50% iEndo-mKate2 EBs in microwell arrays cultured for 1 day. Right, WT-eGFP PGP1 and 50% iEndo-mKate2 EBs cultured for 3 days. **c**, Different seeded proportions of WT-eGFP PGP1 and iEndo-mKate2 hiPSC aggregates in microwell arrays 3 days before suspension culture. Left, 100% iEndo-mKate2. Middle-left, 67% iEndo-mKate2 and 33% WT-eGFP PGP1. Middle-right 33% iEndo-mKate2 and 67% WT-eGFP. Right, 100% WT-eGFP PGP1. **d**, WT-eGFP PGP1 and iEndo-mKate2 EBs 1 day before suspension culture. Left, 100% iEndo-mKate2. Middle-left, 67% iEndo-mKate2 and 33% WT-eGFP PGP1. Middle-right 33% iEndo-mKate2 and 67% WT-eGFP PGP1. Right, 100% WT-eGFP PGP1. Scale bars: 500 μm in **a** (left), **b** (left), and **c**; 100 μm in **a** (right), **b** (right), and **d**.

**Fig. S4.**
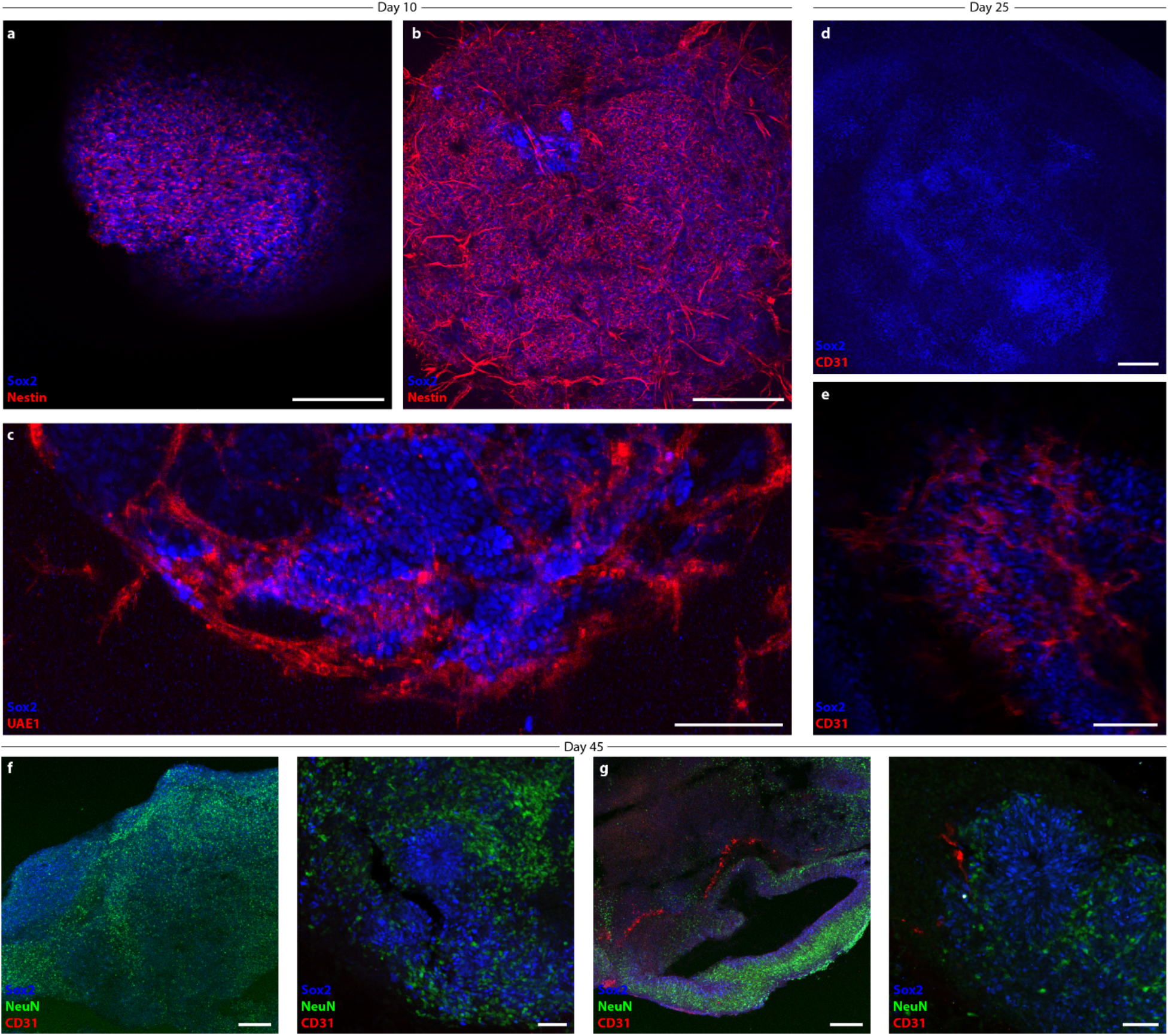
Programmable vascularized cortical organoids. **a**, Immunostaining of Sox2 and nestin in 100% WT PGP1 organoids cultured for 10 days. **b**, Immunostaining of Sox2 and nestin in 67% WT PGP1 and 33% iEndo organoids cultured for 10 days. **c**, Immunostaining for Sox2 and UAE1 for 67% WT and 33% iEndo organoid cultured for 10 days. **d**, Immunostaining of Sox2 and CD31 in 100% WT PGP1 organoids cultured for 25 days. **e**, Immunostaining of Sox2 and CD31 in 67% WT PGP1 and 33% iEndo organoids cultured for 10 days. **f**, Immunostaining of Sox2, NeuN, and CD31 for 100% WT PGP1 organoids cultured for 45 days. **g**, Immunostaining for Sox2, NeuN, and CD31 in a 67% WT PGP1 and 33% iEndo organoids cultured for 45 days. Scale bars: 200 μm in **a**, **b**, **d**, **f** (left), and **g** (left), 100 μm in **c** and **e**, and 50 μm in **f** (right) and **g** (right).

**Fig. S5.**
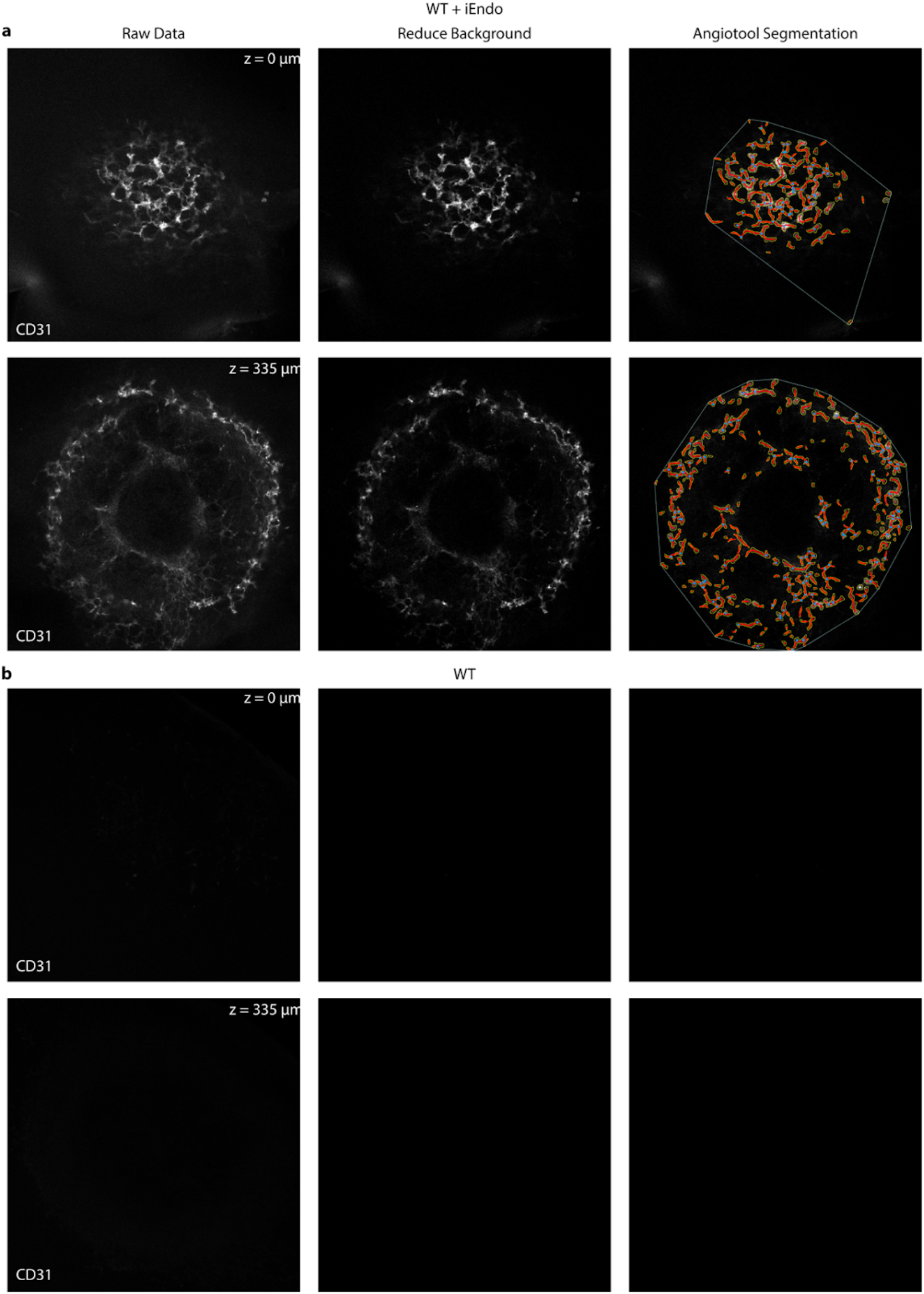
Quantifying vascularization within programmable cortical organoids. Example Angiotool processing workflow for **a**, 67% WT PGP1 and 33% iEndo organoids cultured for 25 days. **b**, 100% WT organoids cultured for 25 days. Left, Individual z sections taken from a confocal z-stack of iDISCO cleared organoids. Middle, Background is eliminated by increasing the minimum threshold on all images. Right, Angiotool analysis of the corresponding z sections. Red lines represent vascular paths, blue dots represent vascular junctions, yellow lines represent the boundaries of the vasculature, and the thin white line marks the calculated vascularized area. Vessel area is calculated as the total area enclosed by the yellow lines.

**Fig. S6.**
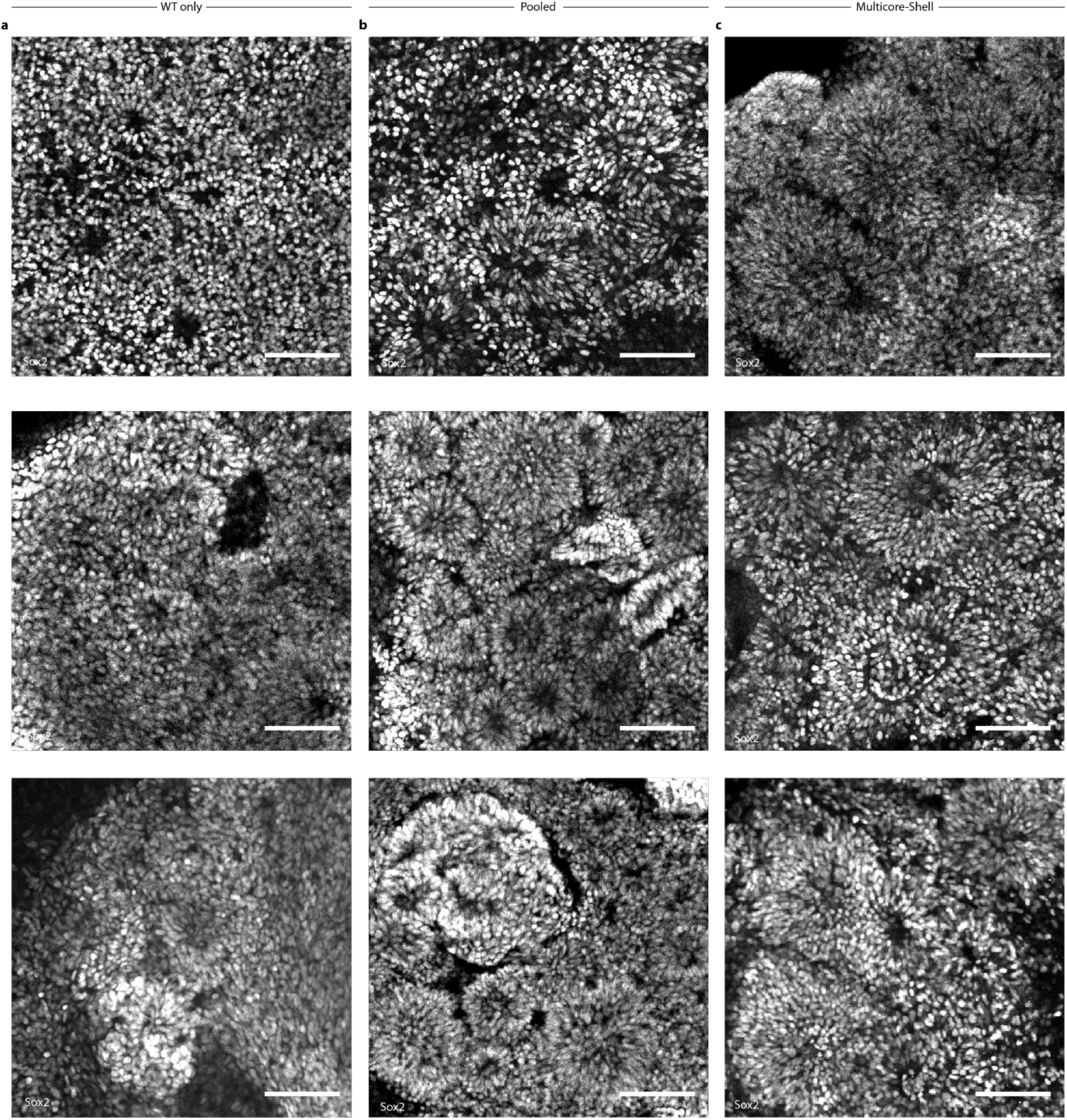
Large ventricular architectures in programmable multicore-shell cortical organoids. **a**, Immunostaining of Sox2 for 100% WT organoids cultured for 10 days. **b**, Immunostaining of Sox2 for randomly pooled organoids cultured for 10 days. **c**, Immunostaining of Sox2 for multicore-shell organoids cultured for 10 days. Scale bars: 100 μm in **a-c**. Cryosection images: **a** (top), **b** (top), **c** (top, middle, bottom). Whole mount images: **a** (middle, bottom), **b** (middle, bottom).

**Fig. S7.**
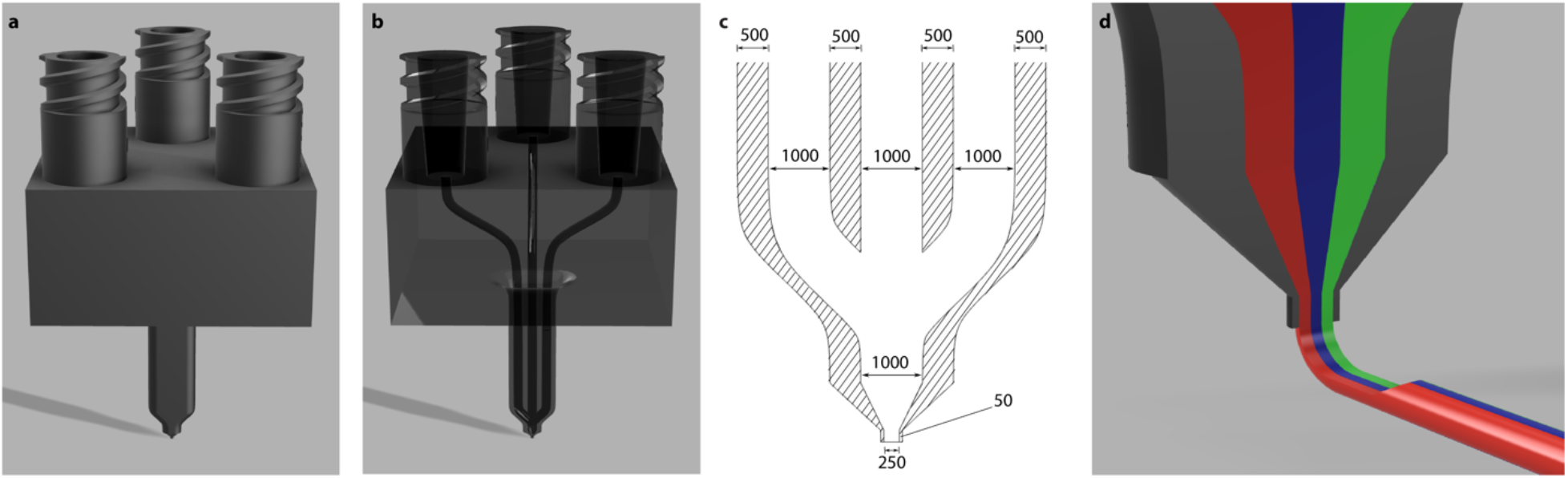
Multimaterial printhead design for creating 3D multicellular tissue architectures. **a**, Perspective view of a 3D CAD model of the triple nozzle. **b**, Perspective view of a translucent 3D CAD model of the triple nozzle visualizing the channels inside the printhead. **c**, Cross-sectional view and dimensions of the printhead tip of the triple nozzle **d**, Cross-sectional view of a 3D CAD model of the triple nozzle and the coextrusion of the three bioinks (red, blue, green). Units in **c** are μm.

**Fig. S8.**
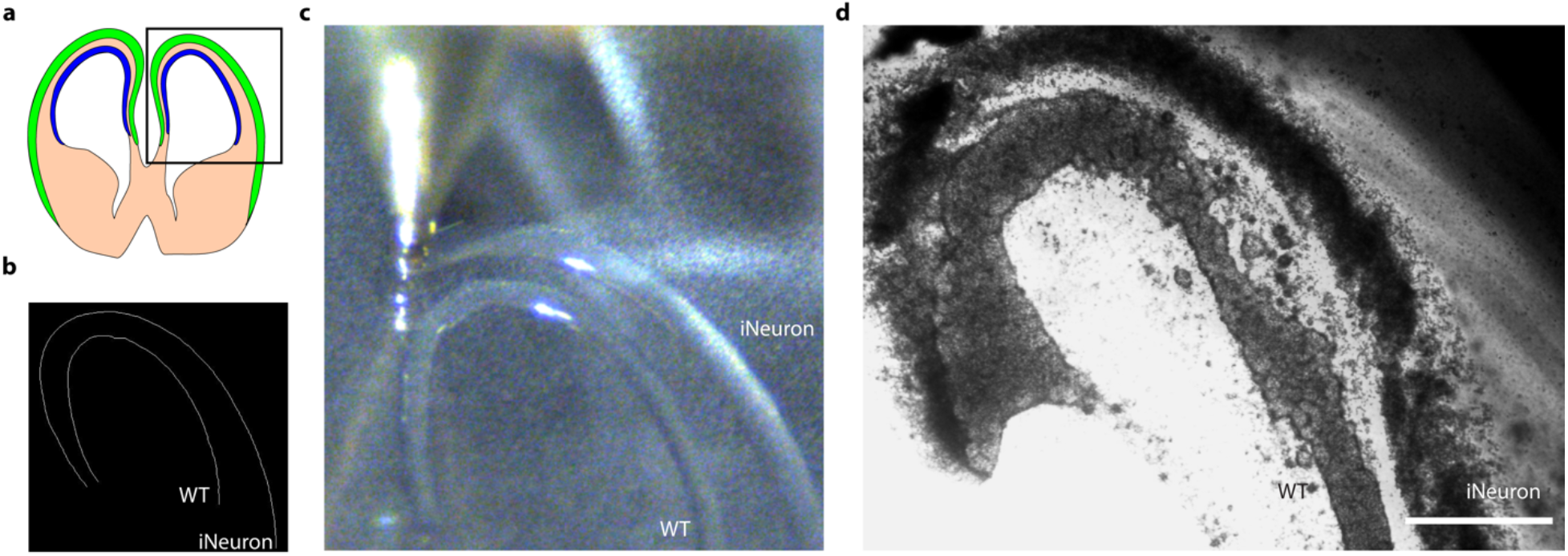
Bioprinting of 3D multicellular tissue architectures resembling a germinal zone and overlying neuronal layer. **a**, Illustration of a developing fetal human brain section in GW11 containing the ventricular zone (shown in blue) and the cortical plate (shown in green). **b**, Extracted printing line path of ventricular zone and neuron dense zone from the brain slice. **c**, Image of nozzle, bioink, and print surface during brain slice printing. **d**, Brightfield image of printed brain slice structure cultured for 5 days. Scale bar: 1 mm in **d**.

**Supplementary Table 1:** Fluorescently conjugated antibodies for flow cytometry.

| Name | Fluorophore | Vendor | Clone | Catalog Number | Dilution |
| --- | --- | --- | --- | --- | --- |
| OCT3/4 | AF647 | BD Biosciences | 40/Oct-3 | 560329 | 1:20 |
| PAX6 | PE | BD Biosciences | O18-1330 | 561552 | 1:20 |
| PAX6 | AF488 | BD Biosciences | O18-1330 | 561664 | 1:20 |
| VE-Cadherin | BV421 | BD Biosciences | 55-7H1 | 565671 | 1:20 |
| MAP2 | PE | MilliporeSigma | AP20 | FCMAB318PE | 1:100 |

**Supplementary Table 2:** Primary immunostaining antibodies.

| Name | Host Animal | Vendor | Catalog Number | Dilution |
| --- | --- | --- | --- | --- |
| CD31 | Mouse | Abcam | ab9498 | 1:200 |
| Nestin | Mouse | Abcam | ab6320 | 1:200 |
| vWF | Mouse | Abcam | ab194405 | 1:200 |
| CTIP2 | Rat | Abcam | ab18465 | 1:200 |
| ZO1 | Rabbit | Abcam | ab96587 | 1:100 |
| NRP1 | Rabbit | Abcam | ab25998 | 1:250 |
| PAX6 | Rabbit | BioLegend | PRB-278P | 1:200 |
| N-Cadherin | Mouse | Cell Signaling Technologies | 14215 | 1:200 |
| VE-Cadherin | Rabbit | Cell Signaling Technologies | 2500S | 1:200 |
| NeuN | Rabbit | Cell Signaling Technologies | 24307 | 1:200 |
| TBR1 | Rabbit | Cell Signaling Technologies | 49661S | 1:200 |
| tRFP | Rabbit | Evrogen | AB233 | 1:200 |
| MAP2 | Mouse | MilliporeSigma | MAB3418 | 1:200 |
| SOX2 | Goat | R&D Systems | AF2018 | 1:200 |
| β-III-Tubulin | Mouse | R&D Systems | MAB1195 | 1:400 |
| OCT3/4 | Rabbit | Santa Cruz Biotechnology | sc-5279 | 1:200 |
| Ulex-Rhodamine | N/A | Vector Laboratories | RL-1062 | 1:200 |

**Supplementary Table 3:**
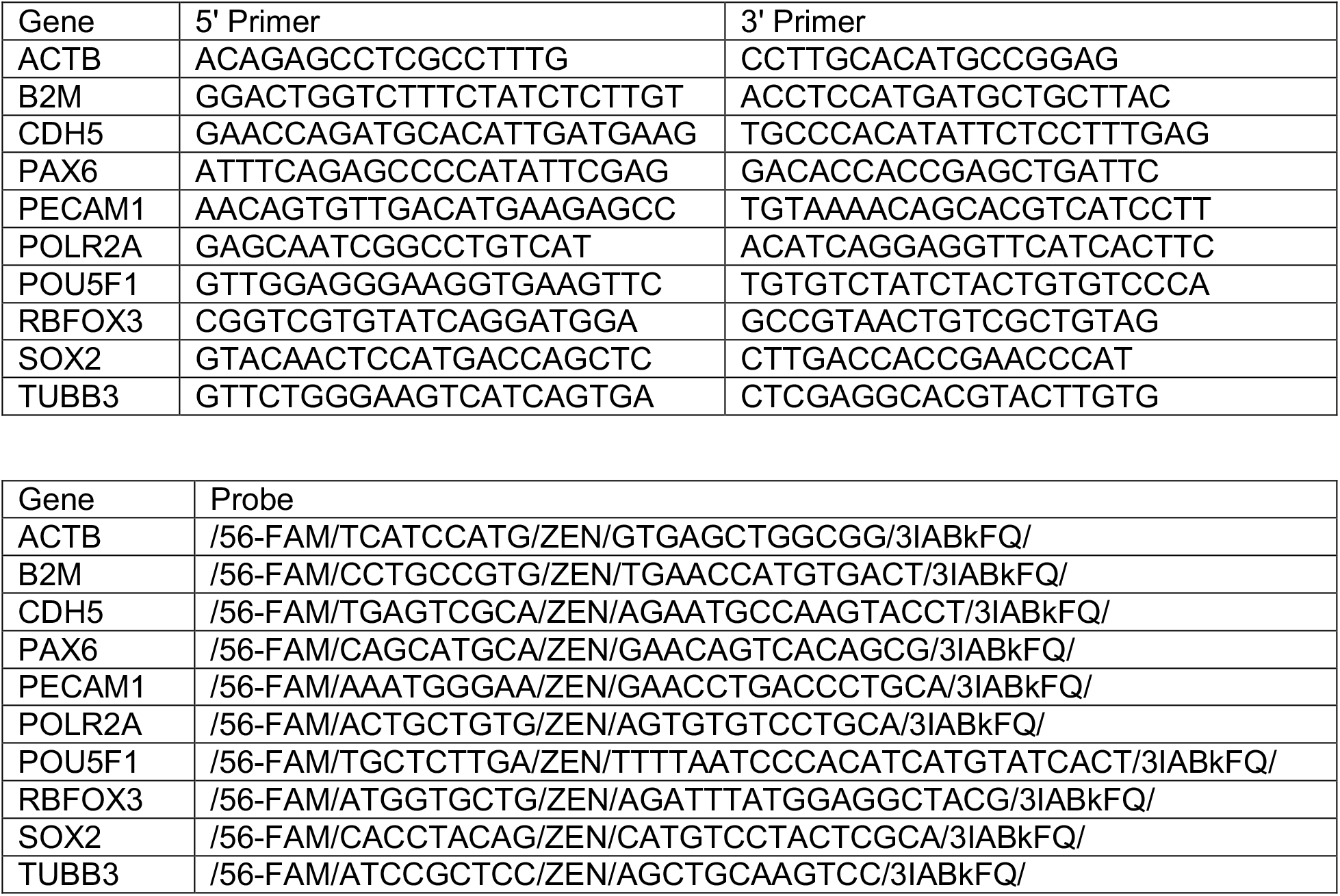
Primers and probes for RT-qPCR.

**Supplementary Video 1 | Cellular origin and evolution of prototypical neurosphere and vascular structures**. Video demonstrates the formation of vascular network and neural rosettes in co-culture over a four-day period on a Matrigel-coated surface in dual-SMAD inhibiting medium. Cells in green are eGFP-expressing WT PGP1 hiPSCs and cells in red are mKate2-expressing iEndo PGP1 cells. Scalebar: 500 μm.

**Supplemental Video 2 | 3D renderings of pervasive vascularization in programmable cortical organoids.** video of confocal z-stack of iDISCO cleared WT and WT + iEndo organoids cultured for 25 days, containing neural stem cell structures marked by Sox2 and vascular structures marked by CD31. Scalebar: 200 μm.

**Supplemental Video 3 | Bioprinting of densely cellular, matrix-free stem cell inks.** Bioprinting of a pseudo-Hilbert curve using a densely cellular, matrix-free hiPSC ink on a transwell membrane.

**Supplemental Video 4 | Bioprinted iNeuron hiPSC filamentary features efficiently differentiate into densely cellular neural filaments.** Confocal z-stack video of printed iNeuron-derived neuron filament in neural induction medium cultured for 6 days. Scalebar: 200 μm.

**Supplemental Video 5 | Multimaterial bioprinting of developmentally inspired multicellular brain tissue architectures.** Bioprinting of layered cortical construct using WT- and iNeuron-bioinks. Video shows the printing of the iNeuron ink alongside a printed WT ink.

## Methods

### Cell culture

PGP1 hiPSCs were utilized for this study (Coriell, #GM23338). Cells were verified for pluripotency by flow-cytometry and cultured between passages 20 and 50. PGP1s were cultured and passaged without antibiotics in mTeSR1 (STEMCELL Technologies, #05850) on tissue culture plates coated with hESC-qualified Matrigel (Corning, #354277) during inducible cell line generation or with growth-factor reduced Matrigel (Corning #354230) for growing hiPSCs prior to differentiation, organoid formation, or bioprinting. For passaging, hiPSCs were washed with phosphate buffered saline (PBS) without calcium and magnesium, dissociated using TrypLE Express (Life Technologies, #12604013), and then seeded at 300k cells/well in a 6-well plate supplemented with 10 μM Y-27632 (Selleck Chemicals, #S1049) for 1 d, and subsequently maintained in mTeSR1 with daily media changes. For cryo-storage, cells were dissociated with TrypLE Express, counted, and resuspended in mFreSR (STEMCELL Technologies, #05854) at a concentration of 10^6^ cells/ml (STEMCELL Technologies, #5854) using a CoolCell LX freezing block (Biocision, #BCS-405) overnight at −80°C, then stored in liquid nitrogen for long-term storage.

### TF-induced clonal hiPSC cell lines

A Gateway compatible, doxycycline-inducible (Tet-On), PiggyBac vector with a puromycin resistance gene was provided by the Church Lab. pDONR221-NEUROG1 (HsCD00040492) from the DNASU^39,40^ repository was cloned into PBAN001 using a Gateway reaction with Gateway LR Clonase II Enzyme mix (Invitrogen, #12538200). The construct was delivered into PGP1 via Super PiggyBac Transposase Expression vector (SBI, #PB210PA-1) by nucleofection at a mass ratio of 1:4 (PBAN-NGN1):(SPB) in 2 μl of total reaction volume, utilizing the Lonza 4D Nucleofector X-Unit (Lonza, #AAF-1002X) with a 20 μl P3 solution kit (Lonza, #V4XP-3032) with 600k PGP1 WT cells in each 20 μl reaction well as per manufacturer’s instructions. After nucleofection, cells in solution were plated on a 6-well tissue culture-treated plate coated with Matrigel in mTeSR1 supplemented with 10uM Y-27632 for one day. Selection by addition of 1 μg/ml puromycin (Gibco, #A1113803) occurred after cells reached 70% confluency and supplemented with 5 μM Y-27632 for one day. Cells were washed once using PBS without calcium or magnesium (Gibco, #14190250) and maintained on mTeSR1 supplemented with 10 μM Y-27632 until colonies were visible.

Confluent polyclonal iNeuron colonies were then dissociated with TrypLE for 7 min, centrifuged at 300 g for 5 min, and resuspended in 500 μl of mTeSR1 supplemented with 10 μM Y-27632 and penicillin-streptomycin (MilliporeSigma, #P4333), strained through a 35 μm cell-strainer (Falcon, #352235) and immediately placed on ice. We used the SH-800s Sony Flow Sorter (100 μm chip nozzle) to sort single cells into Matrigel-coated 96-well plates with each well containing 150 μl of mTeSR1 supplemented with CloneR (STEMCELL Technologies, #05888). Following the manufacturer’s instructions, we maintained the cells until confluent and seeded candidate colonies into two 48-well plate wells, one for validation through induction via addition of 500 ng/ml doxycycline hyclate (MilliporeSigma, #D9891), and another well for further expansion/passaging. iEndo cells, a clonal PGP1 ETV2 isoform 2 line, was constructed in a similar fashion to the iNeurons using the TFome transcription factor library^41^.

To engineer fluorescently labelled hiPSC lines iEndo-mKate2 and eGFP-WT PGP1, the FUGW plasmid (Addgene #14883), which encodes constitutively expressed GFP, was modified to express membrane-bound mKate2 (FUmemKW)^41^. These two plasmids were packaged into lentiviruses as described previously, and FUGW and FUmemKW lentiviruses were transduced into wildtype PGP1 and ETV2 hiPSCs to generate green and red cell lines respectively. hiPSCs were sorted on a FACSAria to obtain a pure population of cells that express the fluorescent proteins uniformly.

### Fluorescent time-lapse imaging during stem cell differentiation

500 μl of cold Matrigel was added to a single well in a 6-well tissue-culture treated plate and allowed to gel by warming to 37°C for 30 min. 150k WT-eGFP PGP1 cells and 150k iEndo-mKate2 PGP1 cells were seeded in mTeSR1 supplemented with Y-27632 and 500 ng/ml doxycycline and cultured for 1 d. On day 2 of culture, the mTeSR1 medium was removed and the plate was gently rinsed with PBS before replacing with neural induction media (NIM), consisting of DMEM/F12 with GlutaMAX (Gibco, #10565018) supplemented with 1:100 N2 supplement (Gibco, #17502048), 1:100 minimal essential media non-essential amino acids (MEM-NEAA), 1 μg/ml heparin (Sigma, H3149), 5 μM SB431542 (BioGems, #3014193), and 100 nM LDN193189 (BioGems, #1066208), with 500 ng/ml doxycycline hyclate added. The plate was transferred to an inverted fluorescence microscope (Zeiss Axio Observer Z1) with an environmental chamber (CO_2_ and temperature controlled) and imaged through a 10× objective onto a Photometrics Evolve electron-multiplying charge-coupled device (EMCCD), tiling over a 4×4 grid every 30 min. Media was changed daily and images were captured over a 4 d period. Tiled images were stitched automatically in Zeiss’ Zen Blue software with a 5% tile-overlap and processed using ImageJ.

### Flow cytometry

For intracellular and surface marker staining, cells were dissociated using Trypsin-EDTA, fixed for 15 min in BD Cytofix (BD, #554714) and washed 3× in PBS with 3% wt/vol bovine serum albumin (BSA). Samples were then aliquoted in a 96-well at 5×10^5^ cells/well, permeabilized in BD Perm/Wash (BD, #554723) solution for 15 min, pelleted via centrifugation at 300 g, and the supernatant was aspirated. Samples were incubated for 45 min in antibodies diluted in BD Perm/Wash at a concentration of 10^7^ cells/ml per dilution in the dark and washed three times in BD Perm/Wash before flow cytometry experiment was performed. Color compensation was performed on each run utilizing BD Anti-Mouse IG (BD, #552843) and fluorescently conjugated antibodies. All flow cytometry measurements were performed on the BD LSR Fortessa cell analyzer. Flow files (.fcs) were processed in FlowJo 10.6.1 (gating strategy included in **Fig. S1**). Plots were generated with Prism 8.4.0. Fluorescently conjugated antibodies utilized in flow cytometry assays are listed in **Supplementary Table 1**.

### HUVEC-hiPSC aggregate cohesion assay

HUVEC-RFP cells (Angio-Proteomie, #cAP-0001RFP), WT-eGFP PGP1 cells, and iEndo-mKate2 PGP1 cells were dissociated from 70%-80% confluency by incubating cells in Gentle Cell Dissociation Reagent (STEMCELL Technologies, #07174) for 12 min at 37°C, then resuspended in DMEM/F12 with HEPES (Gibco, #11330032). Cells were then centrifuged at 250 g for 5 min before resuspending and single-cell filtering in either EB culture media (EBCM) for iEndo-mKate2 cells and WT-eGFP cells, consisting of mTeSR1 supplemented with 4 mg/ml polyvinyl alcohol (PVA, MilliporeSigma, #P8136), or a 1:1 mix of EBCM and EGM-2 (PromoCell, #C-22111) for 1:1 cocultures of HUVEC-RFPs:WT-eGFPs. The PVA stock solution was prepared by fully dissolving PVA in stirred deionized water at 90°C to a stock concentration of 200 mg/ml. Wells of a 24-well AggreWell™ 400 plate (STEMCELL Technologies, #34411) were seeded with either 1:1 iEndo-mKate2:WT-eGFP cells in EBCM with 10 μM Y-27632, or a 1:1 HUVEC-RFP cells:WT-eGFP cells in a 1:1 mix of EBCM and EGM-2, with 10 μM Y-27632. 24 h after plating, media in both conditions was changed to EBCM without Y-27632. 48 h after plating, aggregates were transferred to suspension culture in non-adherent T25 flasks in EBCM on an orbital shaker rotating at 53 RPM. Aggregates were imaged 24 h after plating and 48 h after plating on a Zeiss LSM710 confocal microscope.

### Vascularized cortical organoid culture

Cortical organoid culture was adapted from previously established protocols^12,42,43^. EBs were formed on day –3 by dissociating hiPSC monolayers at 70-80% confluency with EDTA dissociation reagent (EDR) for 12 min at 37°C, 5% CO_2_, which consists of 0.5 mM EDTA in PBS, supplemented with an additional 0.03 M NaCl. Cells were then resuspended in DMEM/F12 with HEPEs and centrifuged at 250 g for 5 min. Wells of a 24-well AggreWell™ 800 plate (STEMCELL Technologies, #34815) were treated with an anti-adhesive coating of 0.2% Pluronic F127 in PBS and seeded with 1.5×10^6^ cells/well using either WT PGP1 hiPSCs or a 1:2 ratio of iEndo:WT PGP1 hiPSCs in EBCM supplemented with 10 μM Y-27632. The day after aggregation (day –2), EBCM media was replaced with fresh EBCM to remove Y-27632 within the AggreWell™ 800 plates. On day –1, EBs were transferred into suspension culture in fresh EBCM in non-adherent T25 flasks on an orbital shaker rotating at 53 rpm.

Neural differentiation began on day 0 (after 1 d of suspension culture). EBs were transferred to neural induction media (NIM) with doxycycline. After 3 d in NIM, organoids were transferred into 80 μl cold collagen/Matrigel gel droplets, formed on dimpled parafilm, which was prepared similarly to a previously established protocol^37^. Final collagen and Matrigel concentrations were 4 mg/ml rat tail collagen type I (Corning, #354249) and 25% Matrigel (Corning). Gel droplets containing organoids were gelled at 37°C for 15 min before being transferred back into NIM in ultra-low adherence 6-well plates (Corning, #3471), which were held stationary. 3 d after implanting organoids into gels (day 6 after the start of neural induction), organoids were transferred onto an orbital shaker rotating at 90 RPM and the media was changed to neural differentiation media 1 (NDM1), consisting of a 1:1 mix of DMEM/F12 with GlutaMAX and Neurobasal media (Gibco, #21103049) supplemented with 1:200 GlutaMAX (Gibco, #35050061), 1:200 MEM-NEAA, 1:200 N2 supplement, 1:100 B27 supplement without vitamin A (Gibco, #12587010), 1:4000 insulin (MilliporeSigma, #I9278), 10 ng/ml VEGF (PeproTech, #100-20), 20 ng/ml EGF (PeproTech, #AF-100-15), 20 ng/ml FGF2 (PeproTech, #100-18B), 50 μM ß-mercaptoethanol (MilliporeSigma, #M6250), and 500 ng/ml doxycycline. 4 d later, half the media was replaced with fresh NDM1. On day 13, after the start of neural induction, a full media changed was performed to replace all media with neural differentiation media 2 (NDM2), consisting of a 1:1 mix of DMEM/F12 with GlutaMAX and Neurobasal media supplemented with 1:200 GlutaMAX, 1:200 MEM-NEAA, 1:200 N2 supplement, 1:100 B27 supplement (Gibco, #17504044), 1:4000 insulin, 10 ng/ml VEGF, 20 ng/ml EGF, 20 ng/ml FGF2, 50 μM ß-mercaptoethanol, and 500 ng/ml doxycycline. Half media changes of NDM2 were performed every 4 d until 25 d after neural induction. 25 d after the start of neural induction, a full media change was performed to replace all media with NDM3, consisting of a 1:1 mix of DMEM/F12 with GlutaMAX and Neurobasal media supplemented with 1:200 GlutaMAX, 1:200 MEM-NEAA, 1:200 N2 supplement, 1:100 B27 supplement with vitamin A (Gibco, #12587010), 1:4000 insulin, 10 ng/ml VEGF, 20 ng/ml BDNF (PeproTech, #450-02), 50 μM ß-mercaptoethanol, and 500 ng/ml doxycycline. Half media changes of NDM3 were performed as needed, every 1-4 days, for the duration of organoid culture. Vascularized cortical organoids were cultured for up to 45 d.

### Multicore-shell cortical organoid culture

Wild type PGP1-iPSCs were dissociated using EDR and seeded into anti-adhesive treated V-bottom 96-well plates (Globe Scientific, #120130) at a density of 1,000 cells/96-well well in EBCM + 10 μM Y-27632 for 12 h to generate organoid cores.

After 12 h, iNeurons were dissociated using EDR and added to each EB at an additional 2,000 cells/96-well well and were allowed to aggregate for 12 h in EBCM with 10 μM Y-27632. Then, iEndo cells were dissociated using EDR and an additional 2,000 cells were added to each EB and allowed to aggregate over 12 h in EBCM + 10 μM Y-27632. 12 h after the final aggregation, media was fully replaced with EBCM. Neural differentiation was initiated 48 h after the start of aggregation by changing the media to NIM within the 96-well plate. 24 h after initiating neural differentiation, organoids were harvested from the 96-well plate and implanted into 80 μl collagen/Matrigel gel droplets. Multicore-shell organoids were then cultured using the vascularized cortical organoid protocol.

### Cryo-sectioning and immunostaining

Organoids were fixed in 4% paraformaldehyde for 30 min and rinsed 3× in PBS. For cryosections, fixed organoids were incubated for 2 d at 4°C in PBS containing 30% wt/vol sucrose and then transferred into a 1:1 solution of Optimal Cutting Temperature compound (OCT) (Tissue-Tek, #4583) and PBS containing 30% wt/vol sucrose for 90 min. Next, the tissue was placed into a cryostat tissue mold, which was subsequently filled with 100% OCT solution and frozen at −20°C on a cryostat Peltier cooler. The tissue was sectioned using 40- to 60-μm slices and transferred onto a Superfrost Plus glass slide (VWR Inc., #48311-703). Sections were stored at −20°C before immunostaining.

For immunostaining of cryosections and whole cortical organoids, tissues were permeabilized for 30 min in PBS containing 0.1% Triton-X, then blocked for >1 h in PBS containing 2% donkey serum. Next, tissue sections were incubated overnight and whole-mount organoids were incubated for 1-3 d in primary antibodies in PBS containing 2% donkey serum. Tissues were rinsed 3× in PBS containing 0.05% Tween-20 (PBST), then incubated in secondary antibodies in PBS with 2% donkey serum for an equal amount of time as primary antibodies. Cell nuclei were labeled with 300 nM of 4′,6-diamidino-2-phenylindole (DAPI) in PBS for 10 min, followed by 3 rinses in PBST. Tissue sections and whole mount organoids were imaged on a Zeiss LSM710 confocal microscope. All primary and secondary antibodies used in immunostaining are listed in **Supplementary Table 2**.

### iDISCO+ tissue clearing

Cortical organoids were cleared and immunolabeled using an adapted version of the iDISCO+ protocol^44^. Briefly, the organoids were dehydrated using a methanol/water gradient over the course of 6 h, then delipidated using a 67% dichloromethane (DCM)/33% methanol solution for 3 h before bleaching in 5% hydrogen peroxide in methanol overnight. Next, they were rehydrated in a reverse methanol/water gradient over the course of 6 h and transferred into PBS. They were then immunolabeled using the same immunolabeling protocol described above and dehydrated a second time using a methanol water gradient over 6 h before further delipidation in 67% DCM/33% methanol for 3 h. The organoids were then rinsed twice in 100% DCM before rehydration over 6 h with a reverse methanol/water gradient. Finally, they were index matched using EasyIndex (LifeCanvas Technologies) and imaged using a Zeiss LSM710 confocal microscope. 3D renderings of cleared cortical organoids were made using the 3Dscript ImageJ plugin^45^.

### Angiotool analysis of vascularized cortical organoids

Confocal *z*-stacks were taken of iDISCO cleared WT only and WT + iEndo organoids. Individual optical sections at *z* = 10.8, 119, 227, 335, and 442 μm within a single organoid were used for Angiotool analysis^46^. Optical sections were preprocessed by raising the LUT threshold to 20 to eliminate background noise. Angiotool analysis was conducted using a vessel diameter of 5 and a small particles filter of 40.

### RNA extraction and RT-qPCR of cortical organoids

Cortical organoids were cultured using protocols described above. RNA extraction was performed utilizing the Ambion PureLink RNA mini kit (Invitrogen, #12183020). Homogenization of single organoids suspended in Matrigel-collagen hydrogel droplet was performed utilizing a hand-held homogenizer (Bel-art, #F65000-0000) with accompanying RNAse-free, DNAse-free single-use pestles (Bel-art, #F65000-0006) with organoids suspended in 600 μl of lysis buffer, provided in the Ambion PureLink Kit. On-column DNAse I (Invitrogen, #AM2222) digestion was also performed during the RNA-extraction process. Extracted RNA samples were validated utilizing an Agilent 2200 TapeStation for RNA integrity number (RIN) score quantification with all samples utilized in this study validated to be above 8.8. cDNA synthesis was performed using the SuperScript IV First-Strand Synthesis System (Invitrogen, #18091050) with Oligo d(T) primer and with the input cDNA for each sample normalized to 10 ng/μl for a total of 110 ng/reaction. cDNA was validated with included kit controls by gel-electrophoresis. RT-qPCR experiments were performed with IDT’s PrimeTime Gene Expression Primer/Probes (ZEN/FAM), utilizing IDT’s PrimeTime Gene Expression Master Mix (IDT, #1055771). Curves were obtained on a BioRad cf96 qPCR machine. Scripts for data processing of Cq values were created in Python and graphs were made utilizing Prism 8.4.0. A list of all primers and probes used is provided in **Supplementary Table 3.**

### Quantification of organoid patterning

Organoids were mounted in EasyIndex Solution and imaged on a Zeiss LSM710 confocal microscope. A custom MATLAB script was written to analyze CellTracker labeled organoids. Each pixel was assigned to one of the three colors used in the CellTracker study based on the fluorescence intensity and pictures were binarized using a maximum entropy thresholding function. Hell *et al.*^47^ give the following formula to account for the focus shift that results from imaging into high refractive index media:

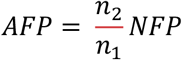

where AFP is the actual focus position (z position of the voxel), NFP is the nominal focus position (imaged z position of the voxel), and *n*_1_ and *n*_2_ are the refractive indexes of air (*n* = 1) and Easy Index (*n* = 1.47), respectively. Easy Index is assumed to have minimal dispersion across visible wavelengths of light. The center position of each organoid is identified, and the distance is calculated between each voxel and the center position for all voxels. Distances were normalized to the radius of each organoid and were summed up to a histogram plot normalized to histogram surface area.

### 3D bioprinting

A triple-material co-flow nozzle (printhead) was designed using Fusion360 (Autodesk Inc.), and exported as a stereolithography (.stl) file. The nozzles were printed using a stereolithography 3D printer (Perfactory Aureus, EnvisionTEC), using HTM140v2 resin (EnvisionTEC) with a layer height of 50 μm and a calibrated power of 700 mW. Printed nozzles were first rinsed and their internal channels were flushed using isopropyl alcohol. They were then dried under a stream of air and further cured under ultraviolet (UV) illumination using an Omnicure lamp (EXFO).

Densely cellular bioinks were created from one 90% confluent T225 flask of PGP1 cells, which were loaded a syringe prior to printing. These iPSCs were first rinsed in PBS without calcium or magnesium and incubated in TrypLE for 7 min at 37°C, 5% CO_2_ and then lifted off and added into 37°C DMEM/F12 with HEPES and centrifuged at 250 g for 5 min. After the supernatant was removed, the cells were resuspended in DMEM/F12 with HEPES, and the suspension was filtered through a 40 μm cell strainer (BD Falcon, #352340) to generate a single cell suspension, which was subsequently pelleted via centrifugation at 250 g for 5 min. The supernatant was aspirated, the remaining cell pellet was resuspended in 250 μl of mTeSR1 containing 10 μM Y-27632, transferred to a 1.6 ml Eppendorf tube, and front loaded into a 1 ml syringe (Covidien Kendall, #8881901014). The 1 ml syringe was centrifuged, with the tip facing upwards, at 750 g for 5 min to form a cell pellet that rests upon the syringe plunger. The supernatant was aspirated inside the syringe by inserting a 1.5 inch-long aspiration nozzle via the tip of the syringe, leaving the pellet intact. Next, the pellet was brought to the tip of the 1 ml syringe by gently manipulating the syringe plunger. The tip of the syringe was then pressed against the back-end of a 250 μl gas-tight glass syringe (Hamilton #81120) without its plunger in place, and the pellet was back-loaded into the glass syringe by gently pressing the 1 ml syringe plunger. Next, the gas-tight plunger was inserted into the glass syringe to bring the rear-loaded pellet up to the tip of the syringe, taking care to avoid introducing air at the plunger-pellet interface. The custom-built printhead was attached to the syringe, which was then mounted on a six-axis motion control stage^9^ fitted with a custom-built syringe pump^10^, for 3D bioprinting.

For bioprinting single-cell filaments, tapered milled metal nozzles with an inner diameter of 50 μm (GPD Global, #10/4794) or 100 μm (GPD Global, #10/4793) were attached to the tip of a glass syringe loaded with the cell pellet. For printing multicellular filaments, a 3D printed triple nozzle was connected to three loaded syringes which were mounted to a single syringe pump that simultaneously drives extrusion of all three inks. Ink extrusion was controlled using an Arduino microcontroller and custom-built stepper motor driver. Printer motion was controlled through using manually written G-code. G-code for the cortical tissue architecture was created from an image taken of a GW11 human brain section^48^. PGP1 hiPSC filaments were printed onto ThinCert transparent 0.4 μm pore sized transwells in a 6-well plate (Greiner Bio-One, #657641). Immediately after printing, the two parts of the gelatin-fibrin pre-gel solution were mixed at a 4:1 ratio and cast overtop the printed filament(s). A combined total of 500 μl of mixed gelatin-fibrin gel was used to encapsulate the printed tissue. 1 ml of EBCM supplemented with 10 μM Y-27632 and 1 U/ml thrombin (MilliporeSigma, #T4648) was added beneath the transwell to keep the cells hydrated. Prints were first incubated for 10 min at room temperature to cross-link the fibrin gel, then transferred in an incubator at 37°C, 5% CO_2_ for 30 min. The media was then removed and replaced with 4 ml/well EBCM supplemented with 10 μM Y-27632, 100 U/ml penicillin-streptomycin, 11.5 KiU/ml aprotinin and 500 ng/ml doxycycline. The next day, media was changed to NIM supplemented with 100 U/ml penicillin-streptomycin and 11.5 KiU/ml aprotinin. Prints were maintained at 37°C, 5% CO_2_ and a full media change was performed every other day until day 6.

After printing, the densely cellular filamentary features were encapsulated in a gelatin-fibrin gel, which was prepared using an modified version of a prior protocol^37^. Briefly, a 15 wt/vol% gelatin solution was produced by adding gelatin powder (MilliporeSigma, #G2500) to PBS without calcium or magnesium and stirring for 12 h at 70°C and adjusting the pH to 7.5 using 1 M NaOH. Part 1 of the gel solution was made by diluting the 15 wt/vol% gelatin 1:1 with mTeSR1 and adding 2.5 mM CaCl_2_, 10 μM Y-27632 and 1 U/mL thrombin for a final 7.5 wt/vol% gelatin mix. Part 2 of the gel solution was produced by dissolving lyophilized bovine blood plasma fibrinogen (MilliporeSigma, #341576) at 37°C in sterile PBS at 50 mg/ml. Both parts of the pre-gel solution were maintained in separate tubes at 37°C before use.

The density of hiPSCs in bioinks was estimated by dispensing 100 μl of bioink from the glass syringe, resuspending the cells in 4 ml of mTeSR1, and counting the number of cells using a cell counter. The density of Oct4+ cells was calculated via multiplying the total number of cells/ml in the bioink by the percentage of cells that were Oct4+, as measured by flow cytometry, using cells obtained from bioink samples.

### CellTracker labeling of hiPSCs

For CellTracker studies of aggregated organoids and printed tissue filaments, CellTracker fluorescent dye was reconstituted in DMSO, then diluted to the working concentration in mTeSR1. The hiPSCs were then incubated with dye for 30 min at 37°C. 2.5 μM of CellTracker™ Green 5-chloromethylfluorescein diacetate CMFDA (Molecular Probes, #C7025) was used to label WT-PGP1 cells, 2.5 μM of CellTracker™ Orange 5-(and-6)-(((4-chloromethyl)benzoyl)amino)tetramethyl-rhodamine CMTMR (Molecular Probes, C2927) was used to label iNeuron cells, and 250 nM of CellTracker™ Deep Red (Molecular Probes, #C34565) was used to label iEndo cells. In triple prints, WT-PGP1 were labeled with 5 μM CellTracker™ Blue 7-amino-4-chloromethylcoumarin CMAC (Molecular Probes, #C2110), iNeuron-PGP1 was labeled with 2.5 μM CellTracker™ Green CMFDA (Molecular Probes, #C7025), iEndo-PGP1 was labeled with 2.5 μM CellTracker™ (Chloromethyl 6-(4(5)-amino-2-carboxyphenyl)-1,2,2,4,8,10,10,11-octamethyl-1,2,10,11-tetrahydrodipyrido[3,2-b: 2,3-i] xanthylium) CMTPX (Molecular Probes, #C34552). Cells were then washed with PBS without calcium or magnesium and dissociated for printing or organoid aggregation. Organoids were fixed 24 h after final aggregation and bioprinted tissues were fixed 4 h after printing in 4% paraformaldehyde for 30 min.

### Statistics and reproducibility

Fig. 2g-I, n=3 independent biological replicates from cells differentiated from different passages; unpaired two-tailed T-test, g, not significant, h, P=7.06×10^−7^, i, P=3.53×10^−5^. Fig. 3j, n=5 optical sections from single whole mount WT and WT+iEndo organoids; unpaired two-tailed T-test, P=4.23×10^−5^. Fig. 3l, n=6 organoids from 3 independent batches. Fig. 4b, n=3 organoids from 3 independent batches. Fig. 5d, n=3 independent bioink batches. Fig. 5e, n=2 (cast control), n=3 (otherwise) independent batches of bioinks.

## Data availability

The datasets generated or analyzed during the current study are available from the corresponding author on reasonable request.

## Acknowledgements

This research was funding by the National Human Genome Research Institute of the NIH under award number RM1HG008525 (M.A.S.-S., J.A.L., J.H., A.N., G.M.C.) and the Vannevar Bush Faculty Fellowship Program (M.A.S.-S., J.A.L), sponsored by the Basic Research Office of the Assistant Secretary for Defense for Research and Engineering through the Office of Naval Research Grant N00014-16-1-2823. A.L. received fellowship support from the Charles Stark Draper Laboratory. The authors also thank T. Ferrante and the Harvard Center for Biological Imaging for microscopy infrastructure and support, S. Uzel, S. Han, and M. Mata for experimental assistance, and J. Coppeta for useful discussions. The authors have submitted a patent application associated with this work.

## Author Contributions

M.A.S.-S. and J.A.L. designed research. M.A.S.-S., J.H., A.L., T.D., L.L.N., and S.D performed research and analyzed data. A.N. and G.M.C. contributed the cell lines used. All authors contributed to manuscript writing.

